# Assembly of a double-stranded RNA synthesizing complex: RNA-DEPENDENT RNA POLYMERASE 2 docks with NUCLEAR RNA POLYMERASE IV at the clamp domain

**DOI:** 10.1101/2020.09.09.290171

**Authors:** Vibhor Mishra, Jasleen Singh, Akihito Fukudome, Feng Wang, Yixiang Zhang, Yuichiro Takagi, Jonathan C Trinidad, Craig S Pikaard

## Abstract

In plants, transcription of selfish genetic elements such as transposons and DNA viruses is suppressed by RNA-directed DNA methylation. This process is guided by 24 nt short-interfering RNAs (siRNAs) whose double-stranded precursors are synthesized by DNA-dependent NUCLEAR RNA POLYMERASE IV (Pol IV) and RNA-DEPENDENT RNA POLYMERASE 2 (RDR2). Pol IV and RDR2 co-immunoprecipitate, and their activities are tightly coupled, yet the basis for their association is unknown. Here, we show that RDR2 stably associates with Pol IV’s largest catalytic subunit, NRPD1 at three sites, all within the clamp module. The clamp is a ubiquitous feature of DNA-dependent RNA polymerases that opens to allow DNA template entry and closes to encase the DNA-RNA hybrid adjacent to the RNA exit channel. The clamp also provides binding sites for polymerase-specific subunits or regulatory proteins, thus RDR2 binding to the Pol IV clamp is consistent with this theme. Within RDR2, the site of interaction with NRPD1 is very near the catalytic center. The locations of the NRPD1-RDR2 contact sites suggest a model in which transcripts emanating from Pol IV’s RNA exit channel align with the template cleft of RDR2, facilitating rapid conversion of terminated Pol IV transcripts into double-stranded RNAs.

**Significance Statement:** Short interfering RNAs (siRNAs) play important roles in gene regulation by inhibiting mRNA translation into proteins or by guiding chromatin modifications that inhibit gene transcription. In plants, transcriptional gene silencing is guided by siRNAs derived from double-stranded (ds) RNAs generated by coupling the activities of DNA-dependent NUCLEAR RNA POLYMERASE IV and RNA-DEPENDENT RNA POLYMERASE 2. We show that the physical basis for Pol IV-RDR2 coupling is RDR2 binding to the clamp domain of Pol IV’s largest subunit. The positions of the protein docking sites suggest that nascent Pol IV transcripts are generated in close proximity to RDR2’s catalytic site, enabling rapid conversion of Pol IV transcripts into dsRNAs.

## Introduction

In eukaryotes, noncoding RNAs guide transcriptional gene silencing to keep transposons, repeated elements and viruses in check, thus defending against genome instability (1–3). These noncoding RNAs include siRNAs and scaffold RNAs to which the siRNAs basepair, an interaction that enables the recruitment of chromatin modifying complexes to the associated chromosomal locus (4–6). In most eukaryotes, noncoding silencing RNAs are dependent on DNA-dependent RNA Polymerase II (Pol II). However, plants evolved two Pol II-derived DNA-dependent RNA polymerases to specialize in noncoding RNA synthesis (7–10), NUCLEAR RNA POLYMERASE IV (Pol IV) and NUCLEAR RNA POLYMERASE V (Pol V) (7), that play non-redundant roles in a process known as RNA-directed DNA methylation (11, 12). Pol IV transcribes chromosomal DNA to generate relatively short transcripts of 25-45 nt (13, 14). An RNA-dependent RNA polymerase, RDR2 (15) then uses the Pol IV transcripts as templates to synthesize complementary strands, yielding double-stranded RNAs (dsRNAs)(16). Resulting dsRNAs are then cleaved by DICER-LIKE 3 (DCL3) to produce short-interfering RNAs (siRNAs) that can be 24 nt or 23 nt in length (16). The 24 nt siRNAs become stably associated with ARGONAUTE 4 (AGO4), or related Argonaute family members, yielding RNA-Induced Silencing Complexes (RISCs) (17–19). RISCs then find their target loci via siRNA basepairing with noncoding scaffold RNAs transcribed by NUCLEAR RNA POLYMERASE V (Pol V) (20, 21). Protein-protein interactions between AGO4 and the C-terminal domain (CTD) of the Pol V largest subunit (22) or between AGO4 and the Pol V-associated protein, SPT5L (23, 24) also contribute to RISC recruitment to sites of Pol V transcription. Once recruited, RISCs facilitate recruitment of the *de novo* DNA methyltransferase, DRM2 (25) as well as histone-modifying activities that act in coordination with DNA methylation (11, 12, 26). Collectively, these activities generate chromatin states that are refractive to promoter-dependent gene transcription, thus accounting for gene silencing.

Using purified Pol IV, RDR2 and DCL3, the siRNA biogenesis phase of the RNA-directed DNA methylation pathway has been recapitulated *in vitro* (16). These experiments revealed that Pol IV and RDR2 enzymatic activities are coupled, with Pol IV’s distinctive termination mechanism somehow facilitating the direct hand-off, or channeling, of transcripts to RDR2 (27). In keeping with their coordinated activities, Pol IV and RDR2 co-immunoprecipitate in *Arabidopsis thaliana* (28, 29) ands *Zea mays* (the maize RDR2 ortholog is MOP1 (30). However, it is not known whether the two enzymes interact directly or via one or more bridging molecules.

Using several independent methods, we show that recombinant RDR2 directly interacts with the largest catalytic subunit of Pol IV, NRPD1. By testing NRPD1 and RDR2 deletion constructs, synthetic peptide arrays, and chemical crosslinking and mass spectroscopy, we identify three intervals within the predicted clamp module of NRPD1 that interact with sequences near the RDR2 catalytic center. Based on homology modeling to related polymerases, we propose that Pol IV-RDR2 docking places the Pol IV RNA exit channel in close proximity to the RDR2 active site, enabling efficient dsRNA synthesis.

## Results

### Recombinant RDR2 has RNA-dependent RNA polymerase activity

Using a baculovirus vector, we expressed in insect cells a recombinant gene that encodes wild-type RDR2 enginered to have a V5 epitope tag and His tag at the N-terminus. A derivative active site mutant (RDR2-asm; Figure 1A) was similarly expressed. Nickel affinity and gel filtration chromatography yielded highly purified RDR2 (Fig 1A, lanes 3 and 5) whose identities were verified by immunoblotting using anti-V5 or anti-RDR2 antibodies (Fig 1A, bottom). Upon analytical gel filtration chromatography, both proteins elute as single peaks with estimated masses of ~140 kD (Fig 1B, see insert), in good agreement with their predicted monomeric masses of ~134 kD.

**Figure 1.**
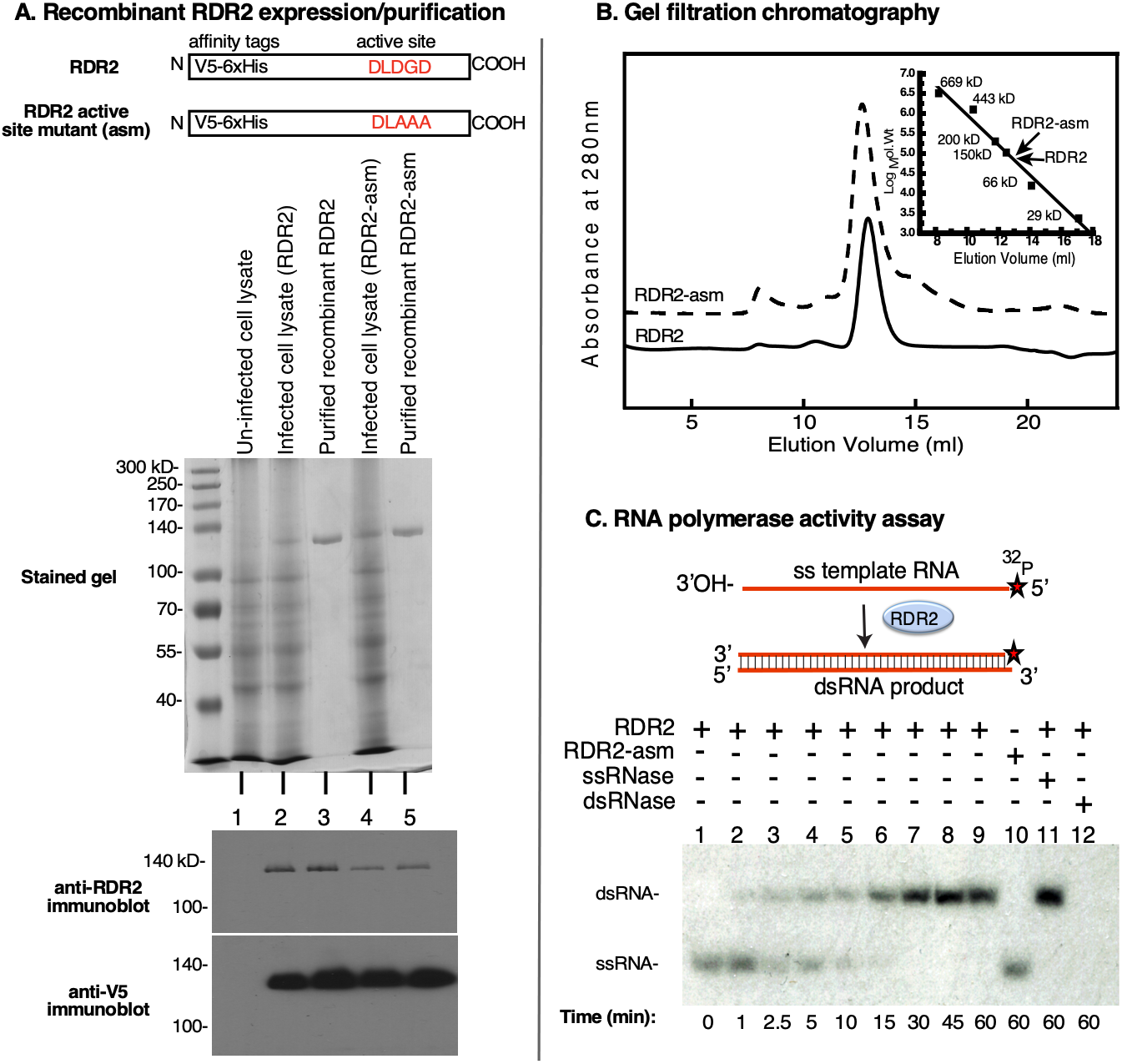
Purification and activity of recombinant RDR2. **A**. The cartoons depict constructs for wild-type RDR2 or active site mutant (asm) that has amino acids of the magnesium binding site changed to alanines. Both proteins have N-terminal V5 epitope tags and 6x His tags. The stained SDS-PAGE gel shows lysates of uninfected High Five™ insect cells, cells infected with the RDR2 or RDR2-asm baculovirus vectors, and the purified proteins. Anti-RDR2 or anti-V5 immunoblots of the samples are shown at the bottom. **B**. Gel filtration chromatography profiles of recombinant RDR2 (solid line) and recombinant RDR2-asm (dashed line). The inset shows a semi-log plot comparing elution volumes of protein mass standards, RDR2 and RDR2-asm. **C**. Conversion of sa ingle-stranded (ss) 37nt RNA, labeled on its 5’ end with ^32^P, into double-stranded (ds) RNA by recombinant RDR2. Lanes 1-9: time-course of conversion of ssRNA into dsRNA. Lane 10: catalytically dead RDR2-asm was substituted for wild-type RDR2. Lanes 11 and 12: reaction products treated with a ssRNA-specific RNase, RNase ONE™ or dsRNA-specific RNase, RNase V1. Reaction products were resolved on a 15% polyacrylamide native gel and visualized by autoradiography.

To test for RNA-dependent RNA polymerase activity, RDR2 and RDR2-asm were provided a ^32^P end-labeled 37 nt single-stranded (ss) RNA template and all four nucleotide triphosphates. Conversion of ssRNA into double-stranded (ds) RNA was then monitored by native gel electrophoresis and autoradiography (Figure 1C). Using RDR2, dsRNA products are detected within 1 minute. In contrast, the RDR2-asm mutant showed no detectable activity, even after 60 minutes (lane 10). Reaction products of RDR2 are resistant to the ssRNA-specific ribonuclease, RNase ONE™ but are sensitive to the dsRNA-specific ribonuclease, RNase V1 (see lanes 11 and 12). Collectively, these results show that recombinant RDR2 can catalyze dsRNA synthesis using an ssRNA template, *in vitro*.

### Pol IV-RDR2 complex reconstitution using recombinant RDR2

Pol IV and RDR2 co-purify and co-immunoprecipitate (co-IP) in both *Arabidopsis thaliana* and maize (28–30). We tested whether Pol IV-RDR2 association can be recapitulated *in vitro* using recombinant RDR2 (Figure 2). For this experiment, we made use of an *A. thaliana* transgenic line in which NRPD1 bearing a FLAG epitope tag at its C-terminus (NRPD1-FLAG) rescues an *nrpd1-3* null mutant (8). This allows Pol IV to be affinity purified using anti-FLAG resin (see cartoon of Figure 2A). Using polyclonal antibodies recognizing native NRPD1 or native RDR2, both proteins are detected upon NRPD1-FLAG IP (Figure 2A lane 2), in agreement with prior studies (29). If NRPD1-FLAG IP is used to affinity-capture Pol IV from a *rdr2* null mutant background, RDR2 is not detected, as expected (lane 3). RDR2 is also not detected in Pol V or Pol II fractions affinity purified by virtue of FLAG-tagged NRPE1 (the largest subunit of Pol V) or NRPB2 (the second-largest subunit of Pol II)(lanes 4 and 5) (8), in keeping with RDR2’s specific association with Pol IV.

**Figure 2.**
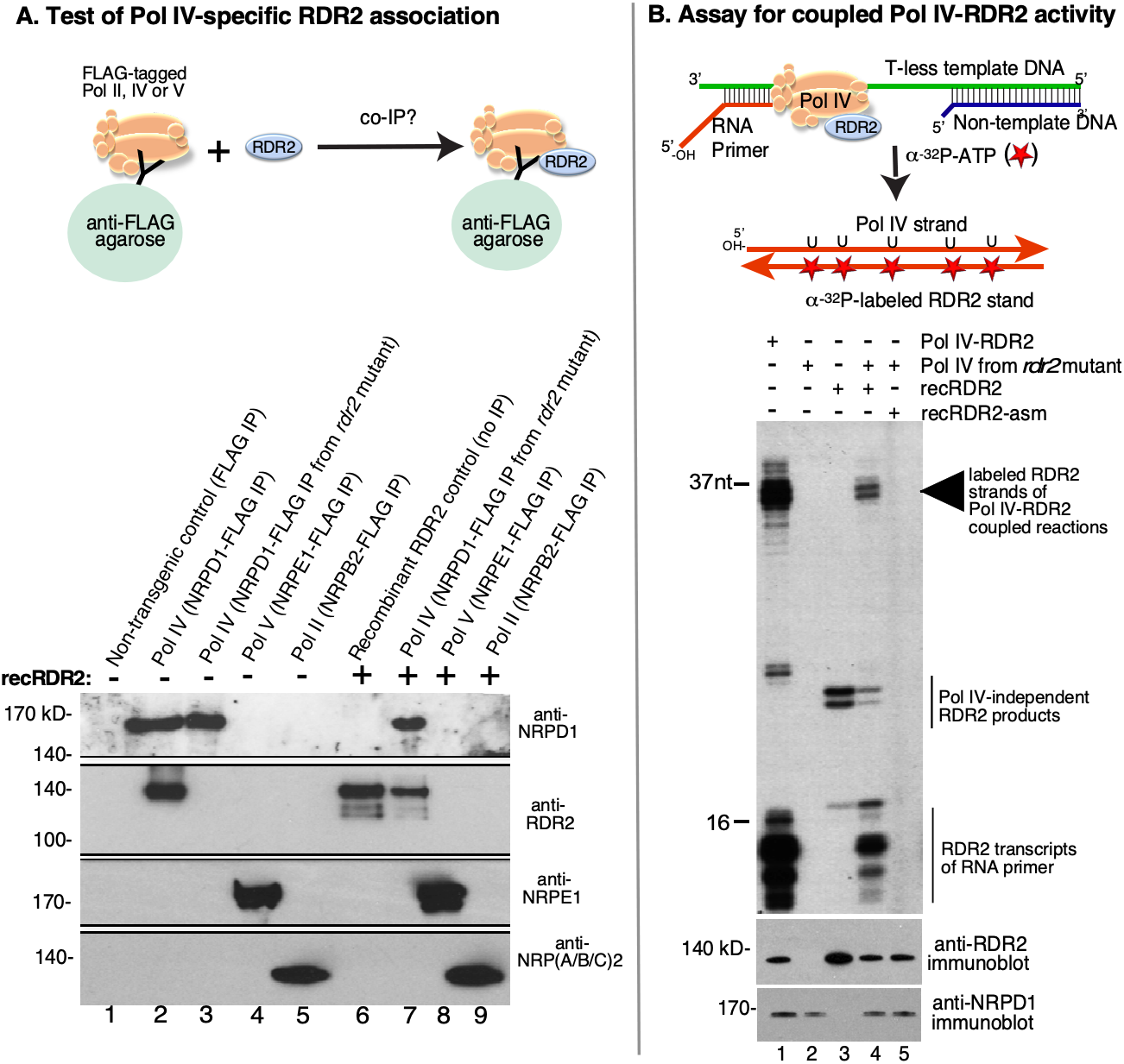
Reconstitution of a functional Pol IV-RDR2 complex. **A**. Recombinant RDR2 stably associates with Pol IV. Pol IV, Pol V, and Pol II assembled using FLAG-tagged NRPD1, NRPE1, or NRPB2, respectively, were immunoprecipitated (IPed) from transgenic plants using anti-FLAG resin. In lane 1, anti-FLAG IP of a non-transgenic plant lysate serves as a negative control. In lane 2, Pol IV was IPed from plants that are wild-type for RDR2. In lanes 3 and 7, Pol IV IP ed from an *rdr2* null mutant. In lanes 7-9, recombinant RDR2 was mixed with the indicated Pol IV, Pol V or Pol II fractions prior to anti-FLAG IP. Lane 6 shows recombinant RDR2 as a positive control. Proteins resolved by SDS-PAGE were subjected to immunoblotting using anti-RDR2, anti-NRPD1, anti-NRPE1 or an antibody that recognizes a peptide sequence of the Pol II subunit, NRPB2 that is also present in the NRPA2 and NRPC2 subunits of Pols I and III, respectively. **B**. Pol IV-RDR2 assembled using recombinant RDR2 carries out the coupled Pol IV-RDR2 reaction, generateing dsRNA from a DNA template. The cartoon illustrates the use of a T-less DNA template and RNA labeling with α-^32^P-ATP to specifically label second RNA strands synthesized by RDR2 (Singh et al., 2019). Lane 1 shows the activity of Pol IV-RDR2 complexes isolated from plants expressing wild-type RDR2. Lane 2 tests the activity of Pol IV purified from the *rdr2* null mutant background. Lane 3 tests the activity of recombinant RDR2 alone. In lanes 4 and 5, Pol IV-RDR2 complexes were reconstituted using recombinant RDR2 or catalytically dead RDR2-asm and tested for activity. The bottom panel shows immunoblots to detect NRPD1 or RDR2 present in the reactions.

Pol IV, Pol V and Pol II were next IPed following incubation with recombinant RDR2 (Figure 2A, lanes 7-9). Importantly, Pol IV purified from the *rdr2* null mutant interacts with recombinant RDR2 such that both proteins co-IP (Figure 2A, lane 7). By contrast, Pols II or V do not pull down RDR2 (lanes 8 and 9).

We next tested whether the Pol IV-RDR2 complexes assembled using recombinant RDR2 can carry out the coupled enzymatic reactions that generate dsRNA from a DNA template strand (16). In this assay, diagrammed in Figure 2B, Pol IV transcription is initiated using an RNA primer hybridized to a T-less (no thymidines) 51 nt DNA template strand. When the elongating Pol IV encounters a 28 nt non-template strand of DNA, basepaired to the template strand, Pol IV transcribes only ~12-16 nt into the double-stranded region and then terminates, yielding transcripts of ~34-37 nt (16). Importantly, Pol IV termination in this manner is required for RDR2 to engage the Pol IV transcript and synthesize the complementary RNA strand (16). Because no thymidines are present in the DNA template, α-^32^P-ATP is not incorporated into the initial Pol IV transcript. However, uracils incorporated into the Pol IV transcripts template the incorporation of α-^32^P-ATP into the RNA second strands that are synthesized by RDR2. Thus α-^32^P-ATP incorporation into RNAs of ~34-37 nt is indicative of coupled Pol IV-RDR2 transcription (16).

Lane 1 of Figure 2B shows reaction products generated by Pol IV associated with wild-type RDR2 (see immunblots at bottom), with prominent 34-37 nt body-labeled RDR2 transcripts being readily apparent. Pol IV purified from an *rdr2* null mutant does not generate these transcripts (lane 2). The ladder of short RNA transcripts near the bottom of lane 1 are RDR2 transcripts generated using the 16 nt RNA primer as a template (29). Haag et al. showed that RDR2 associated with Pol IV generates these short primer transcripts but RDR2 isolated from a *pol iv* mutant does not (29). In keeping with these observations, recombinant RDR2 does not generate the short primer transcripts (Figure 2B, lane 3), although longer transcripts of ~22-23 nt are observed. Adding recombinant RDR2 to the Pol IV fraction reconstitutes the Pol IV-RDR2 coupled reaction (lane 4), generating 34-37 nt transcript like native Pol IV-RDR2 (compare lanes 4 and 1). Short RDR2 transcripts derived from the RNA primer are also observed for the reconstituted Pol IV-RDR2 complex (lane 4). Importantly, no products are observed in reactions in which Pol IV-RDR2 complexes were reconstituted using RDR2-asm (lane 5).

### RDR2 physically interacts with the largest catalytic subunit of Pol IV

Having demonstrated that recombinant RDR2 associates with Pol IV to form a functional complex capable of dsRNA synthesis, we sought to identify the basis for Pol IV-RDR2 association. As an initial test, we performed a far-western blot in which FLAG-tagged Pol II, Pol IV and Pol V were affinity-purified, subjected to SDS-PAGE, electroblotted to nitrocellulose, and incubated with recombinant V5-tagged RDR2. Filters were washed extensively then incubated with anti-V5 antibody conjugated to horseradish peroxidase (HRP). Development of the blot, by chemiluminescent detection of HRP activity, revealed a single band of ~165 kDa in the Pol IV fraction, consistent with the size of the NRPD1 subunit (Figure 3A, lane 4).

**Figure 3.**
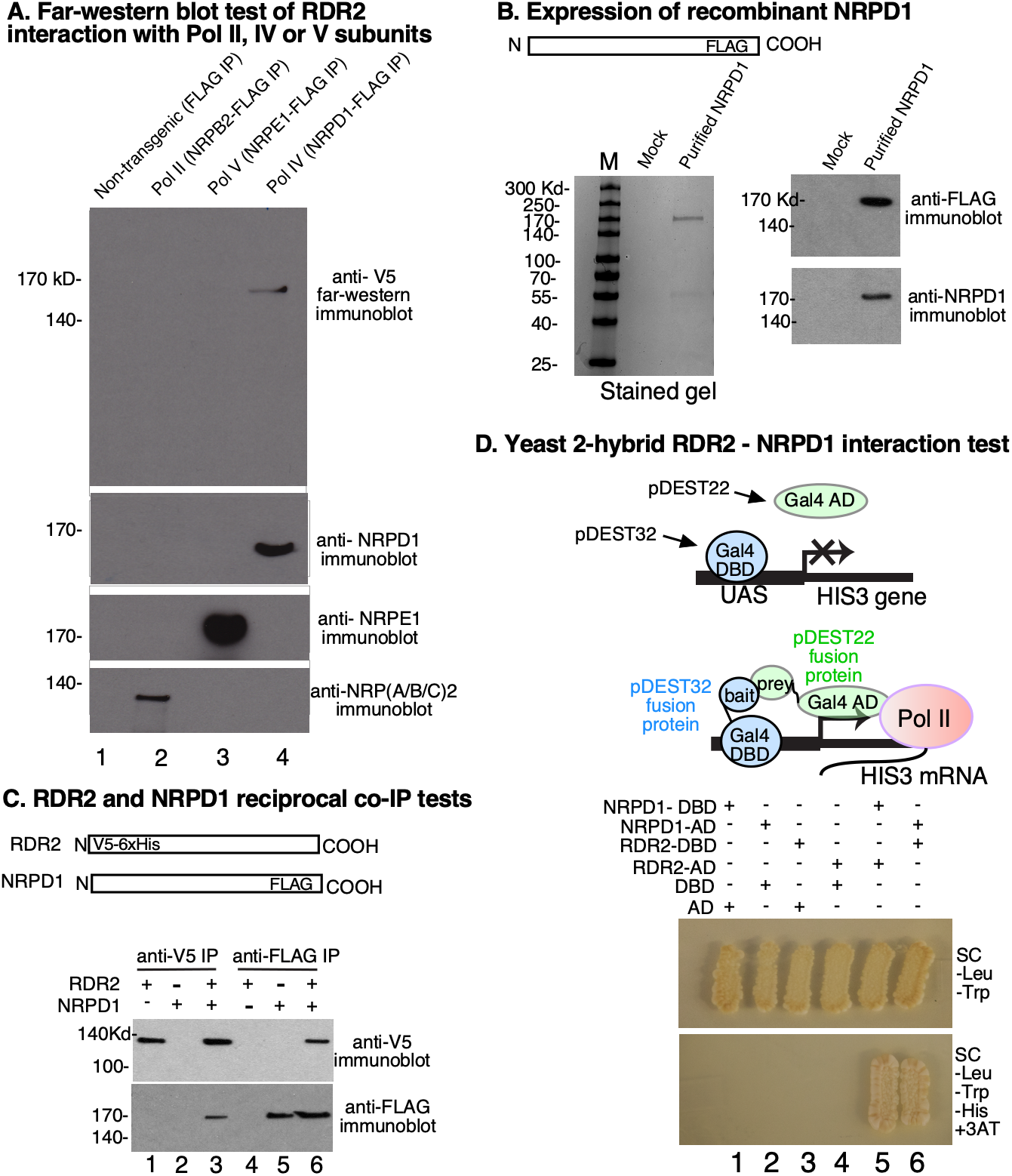
RDR2 interacts with the Pol IV largest subunit, NRPD1. **A**. Far-western blot test for RDR2 interacting proteins. Pols II, V and IV assembled using FLAG-tagged NRPB2, NRPE1 or NRPD1, respectively were IPed using anti-FLAG resin. Following SDS-PAGE and electroblotting, the filter was incubated with recombinant V5-tagged RDR2. After washing, the filter was incubated with anti-V5 antibody to detect immobilized RDR2. Immunoblots at the bottom of the panel detect NRPD1 (Pol IV), NRPE1 (Pol V) or NRPB2. **B**. Expression of recombinant NRPD1. Shown at left is a Coomassie-stained SDS-PAGE gel comparing purified recombinant NRPD1 expressed in High Five™ insect cells to mock-infected cells subjected to the same purification protocol. At right are anti-FLAG and anti-NRPD1 immunoblots of these fractions. **C**. Recombinant NRPD1 and recombinant RDR2 co-immunoprecipitate. V5-tagged RDR2 and FLAG tagged NRPD1 were incubated alone or together, as indicated, then IPed using anti-V5 or anti-FLAG resins. IPed fractions were then subjected to SDS-PAGE and immunoblotting using anti-V5 or anti-FLAG antibodies. **D**. NRPD1 and RDR2 interact in reciprocal yeast two-hybrid interaction tests. The cartoon depicts the experiment, using proteins fused to the Gal4 DNA binding domain (DBD) as bait and proteins fused to the Gal4 activation domain (AD) as prey. Upon co-transformation of the fusion constructs into yeast, the yeast are able to grow on media lacking tryptophan and leucine (Top panel). Interaction between the indicated fusion proteins is demonstrated by growth on media lacking tryptophan, leucine, histidine, and with addition of 3-amino-1,2,3-trizole (3AT) (bottom panel). UAS = upstream activating sequence that GAL4 binds.

To test whether RDR2 can interact with NRPD1, we designed a synthetic transgene encoding NRPD1 fused at its C-terminus to a FLAG epitope tag, expressed the protein in insect cells using a baculovirus vector, and purified the protein to near homogeneity (Figure 3B). Upon incubation with V5-tagged RDR2, the proteins co-IP, using either anti-V5 or anti-FLAG resin, indicating that NRPD1 and RDR2 do, indeed interact (Fig 3C, lanes 3 and 6).

As a third test of RDR2’s ability to interact with NRPD1, we performed reciprocal yeast two-hybrid interaction experiments (Figure 3D) using NRPD1 or RDR2 as either bait (when fused to the Gal4 DNA binding domain, DBD of pDEST32) or prey (when fused to the Gal4 activation domain, AD of pDEST22). Either combination of NRPD1 and RDR2, as bait or prey, resulted in HIS3 expression, enabling colony growth on media lacking histidine and containing 25 mM 3 amino-1,2,3-triazole (3AT) to increase the stringency of HIS selection (Fig 3D, lanes 5 and 6).

### The N-terminal region of NRPD1 interacts with RDR2

Full-length NRPD1 is 1453 amino acids in length. To search for regions that interact with RDR2, we engineered six recombinant expression vectors, each encoding ~33 kDa portions of NRPD1 (Figure S1A), each overlapping its neighbors by ~5 kDa, and each having a C-terminal FLAG epitope tag. The polypeptides were generated *in vitro* using a bacterial transcription-translation expression system (Figure S1A) then incubated with V5-tagged recombinant RDR2 (see cartoon of Figure 4A). Following anti-V5 IP, proteins were subjected to SDS-PAGE and immunoblotting, with anti-V5 antibody used to detect RDR2 and anti-FLAG antibody used to detect NRPD1 polypeptides. The polypeptide corresponding to the amino terminus of NRPD1, amino acids 1-300, co-IPed with RDR2 (Fig 4A, lane 2). The partially overlapping polypeptide corresponding to amino acids 255-555 did not coIP with RDR2 (Fig 4A, lane 3) suggesting that RDR2 interaction likely occurs within NRPD1’s first 255 amino acids.

**Figure 4.**
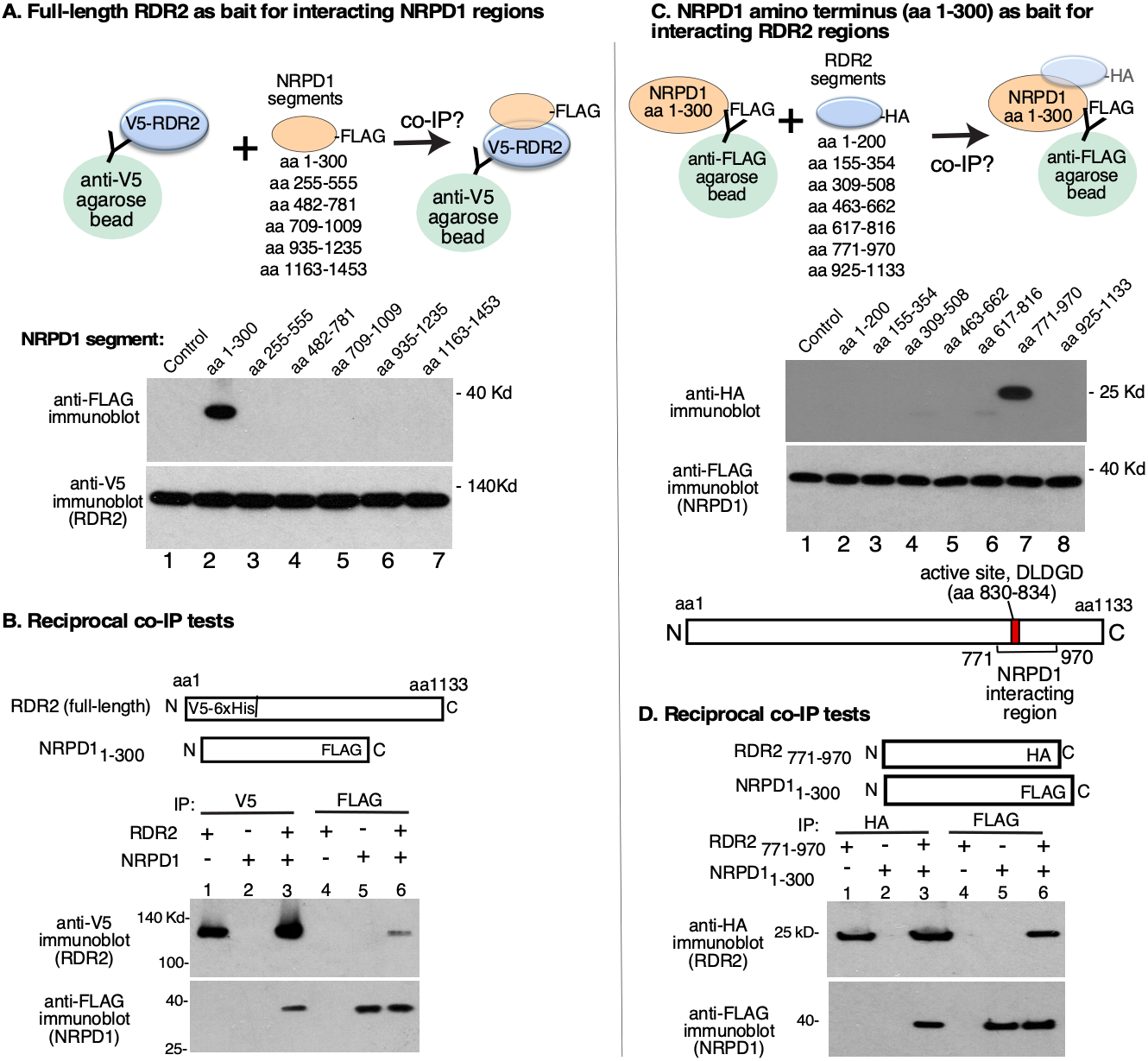
Mapping interacting subdomains of NRPD1 and RDR2. **A**. Testing NRPD1 amino acid intervals for interaction with RDR2. Six overlapping polypeptides that collectively represent the 1453 amino acids of full-length NRPD1 were designed, each having a C-terminal FLAG tag. The polypeptides were then expressed *in vitro* using a cell-free transcription-translation system (see Figure S1A) and incubated with V5-tagged recombinant RDR2. Anti-V5 immunoprecipitation was then used to pull down RDR2 and any associated proteins, as depicted in the cartoon. IPed proteins were resolved by SDS-PAGE and subjected to immunoblotting using anti-FLAG and anti-V5 antibodies. Immunoblots were probed with anti-FLAG and anti-V5. **B**. NRPD1_1-300_ and RDR2 co-IP. Recombinant, FLAG-tagged NRPD1_1-300_ expressed in *E. coli* (see Figure S1B) was incubated with recombinant V5-tagged RDR2 (lanes 3 and 6), then IPed using either anti-FLAG or anti-V5 resin. In lanes 1-2 and 4-5, the individual proteins were IPed as controls. **C**. Testing RDR2 amino acid intervals for interaction with NRPD1_1-300_. Seven overlapping polypeptides of ~22 kDa that collectively represent the amino acid sequence of full-length RDR2 (1133 amino acids) were designed, each having a C-terminal HA epitope tag. The recombinant RDR2 polypeptides were then expressed *in vitro* using a cell-free transcription-translation system (see Figure S2A). The RDR2 polypeptides were then incubated with NRPD1_1-300_ followed by anti-FLAG IP to pull down NRPD1_1-300_ and any associated proteins, as depicted in the cartoon at top. IPed proteins were resolved by SDS-PAGE and subjected to immunoblotting using anti-FLAG and anti-HA antibodies. Immunoblots were probed with anti-HA and anti-FLAG. The diagram at bottom shows that RDR2_771-970_ includes the enzyme’s active site. **D**. Recombinant NRPD1_1-300_ and recombinant RDR2_771-970_, each expressed in *E. coli* (see Figures S1B and S2B) were incubated together (lanes 3 and 6) or alone (lanes 1-2, 4-5), then subjected to anti-HA or anti-FLAG IP. IPed proteins were resolved by SDS-PAGE and subjected to immunoblotting using anti-FLAG or anti-HA antibodies.

To further test the ability of the NRPD1_1-300_ polypeptide to interact with RDR2, we engineered a recombinant gene that expresses the NRPD1_1-300_ polypeptide in *E. coli* and affinity purified the polypeptide by virtue of its C-terminal FLAG tag (Fig S1B). Following incubation with V5-tagged RDR2, reactions were then IPed using anti-V5 or anti-FLAG resin. IPed proteins were then subjected to SDS-PAGE and immunoblotting, using anti-V5 antibody to detect RDR2 and anti-FLAG antibody to detect NRPD1_1-300_ (Figure 4B). In each test, RDR2 and NRPD1_1-300_ co-IPed (see Figure 4B, lanes 3 and 6).

### A site flanking the RDR2 active site interacts with NRPD1

To identify regions of RDR2 that interact with NRPD1, we engineered constructs encoding seven overlapping polypeptides of ~25 kDa, each bearing an HA epitope tag, and expressed the polypeptides *in vitro* using a bacterial transcription-translation system (Figure S2A). Following incubation with FLAG-tagged NRPD1_1-300_, IP was conducted using anti-FLAG resin and affinity-captured proteins were subjected to SDS-PAGE and immunoblotting using anti-HA or anti-FLAG antibodies (Figure 4C). The polypeptide corresponding to RDR2 amino acids 771-970, which includes the conserved aspartate triad of the active site, co-IPs with NRPD1_1-300_ (Fig 4C, lane 7). The partially overlapping polypeptides corresponding to amino acids 617-816 or 925-1133 did not co-IP with NRPD1_1-300_. Collectively, these results suggest that RDR2 sequences between amino acids between 816 and 925 interact with NRPD1.

To confirm the ability of RDR2_771-970_ to interact with FLAG-tagged NRPD1_1-300_, we expressed RDR2_771-970_, fused to an HA epitope tag, in *E. coli* and affinity-purified the protein using anti-HA resin (Figure S2B). We then incubated the protein with FLAG-tagged NRPD1_1-300_ and performed IP using anti-FLAG or anti-HA resin. These tests showed that RDR2_771-970_ and NRPD1_1-300_ co-IP, regardless of which partner is affinity captured (Figure 4D, lanes 3 and 6).

### Fine mapping Pol IV-RDR2 interaction sites using peptide arrays

Having identified regions of NRPD1 and RDR2 that interact, we next searched for peptides within these regions that might account for the interactions. Twenty peptides that collectively comprise the amino acids of RDR2_771-970_ and thirty peptides that comprise the NRPD1_1-300_ sequence were synthesized and arrayed by dot blotting onto nitrocellulose (Figures 5A and B). Peptides were typically 15 amino acids in length and overlapped their neighbors by 5 amino acids. The RDR2 peptide array was incubated with NRPD1_1-300_-FLAG and the NRPD1 peptide array was incubated with RDR2_771-970_-HA. Filters were washed to remove unbound probe proteins and then incubated with HRP-conjugated antibodies recognizing the HA or FLAG tags of the NRPD1_1-300_ or RDR2_771-970_ polypeptides. Filters were again washed, then assayed for HRP-catalyzed chemiluminescence to screen for the presence of immobilized probe polypeptides (see cartoon of Figure 5A).

**Figure 5.**
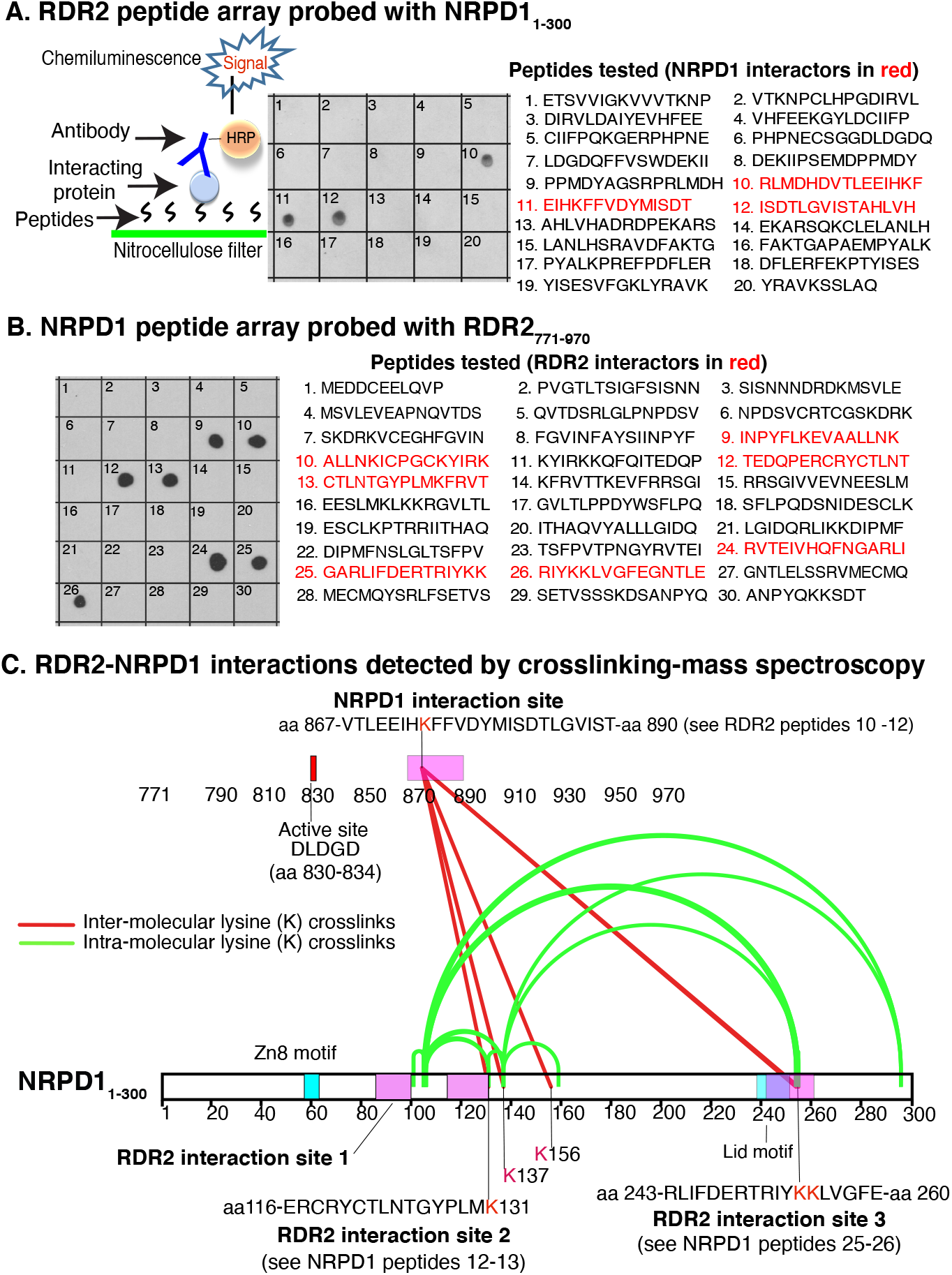
Interacting peptides of NRPD1 and RDR2. A. Peptides of RDR2_771-970_ that interact with NRPD1_1-300_. Twenty overlapping 15-amino acid peptides were dot-blotted onto nitrocellulose and incubated with NRPD1_1-300_-FLAG. After washing, the filter was probed with anti-FLAG antibody conjugated to horseradish peroxidase (HRP). HRP activity was then detected by chemiluminescence. B. Peptides of NRPD1_1-300_ that interact with RDR2_771-970_.Thirty overlapping 15-amino acid peptides were dot-blotted onto nitrocellulose and incubated with RDR2_771-970_-HA. After washing, the filter was probed with anti-HA antibody conjugated to horseradish peroxidase (HRP). HRP activity was then detected by chemiluminescence. C. RDR2 -NRPD1 interactions detected by crosslinking and mass spectroscopy. NRPD1_1-300_ and RDR2_771-970_ were incubated together with the chemical crosslinker BS3 then subjected to SDS-PAGE to resolve crosslinked from non-crosslinked polypeptides. A summary of the mass spectroscopy results for the gel-resolved complexes are shown. Crosslinks between lysines of the two proteins are shown as red lines. Crosslinks between lysines of NRPD1 are shown as green arcs. Interacting peptide regions of the two proteins are depicted as pink rectangles. The Zn8 and Lid elements of NRPD1 are shown as blue rectangles. The overlap between the Lid and RDR2 interaction site 3 is depicted in purple. The data are available via ProteomeXchange via the identifier PXD020170.

Three contiguous peptides of RDR2, peptides 10-12, interact with NRPD1_1-300_ (Figure 5A). If shared sequences of the non-interacting, overlapping peptides 9 and 13 are excluded, the sequence ^866^DVTLEEIHKFFVDYMISDTLGVIST^890^ emerges as the candidate NRPD1 contact region. This sequence is close to the active center, beginning 32 amino acids downstream from the ^830^DLDGD^834^ motif that coordinates a magnesium ion at the site of phosphodiester bond formation (*see* upper diagram of Figure 5C).

In the case of NRPD1, three RDR2-interacting intervals were detected, defined by peptides 9-10, 12-13 and 24-26 (Figure 5B). Omitting shared amino acid sequences of non-interacting adjacent peptides, the sequences ^86^LKEVAALLNKICPGC^100^, ^116^ERCRYCTLNTGYPLM^130^ and ^236^VHQFNGARLIFDERTRIYKKLVGFE^260^ are deduced to contain RDR2 interaction motifs.

### Mapping Pol IV-RDR2 interaction sites by chemical crosslinking and mass spectroscopy

As an independent means of identifying interacting peptides of NRPD1 and RDR2, we incubated NRPD1_1-300_ and RDR2_771-970_ with the lysine crosslinker bis-sulfosuccinimidyl suberate (BS3). We then used SDS-PAGE to gel-purify crosslinked complexes, performed digestion with trypsin and chymotrypsin, and identified resulting peptides by mass spectroscopy (Figure 5C). Crosslinked peptides of each protein were detected, including intra- and inter-protein crosslinks. Crosslinks between RDR2 and NRPD1 occurred between lysine 874 (K874) of RDR2_771-970_ and lysines K131, K137, K156, K254 or K255 of NRPD1_1-300_ (Fig 5C, red lines). Importantly, lysine 874 of RDR2 is present within peptides 10 and 11 of the peptide array (see Figure 5A) and is the only lysine within NRPD1-interacting peptides 10-12. This region of RDR2 is depicted as a pink rectangle in the diagram of Figure 5C. Likewise, two of the three RDR2-interacting regions of NRPD1 previously defined in the peptide array experiment included lysines that were crosslinked to RDR2, specifically K131, which is present in peptides 12 and 13, and K254 and K255, present within peptides 25 and 26 (Figures 5C and 5B). Moreover, crosslinks formed between lysines within NRPD1_1-300_ (green arcs in Figure 5C) indicate that RDR2 interacting region 3, which is seemingly far apart from interacting regions 1 and 2, must actually be in close proximity to these regions in the folded polypeptide because the distance given that the reactive groups of the BS3 crosslinker is only 11.4 Å. Collectively, these results suggests that NRPD1_1-300_ peptides 9-10, 12-13 and 24-26 may comprise a RDR2-binding surface.

### Pol IV-RDR2 interaction sites are conserved in plants

The *A. thaliana* genome encodes six RNA-dependent RNA polymerases (RdRPs), but only RDR2 interacts with Pol IV. The NRPD1 interaction site within RDR2 is deduced to involve the sequence DVTLEEIHKFFVDYMISDTLGVIST, as discussed above. This sequence has little similarity to the corresponding sequences of RDRs 1,3,4,5 or 6 (Figure 6A). By contrast, the nearby active site region is highly conserved, especially between RDR2 and its closest paralogs, RDR1 and RDR6. The conservation at the active site extends to *Neurospora crassa*,QDE-1, for which structural information is available (31)(see below).

**Figure 6.**
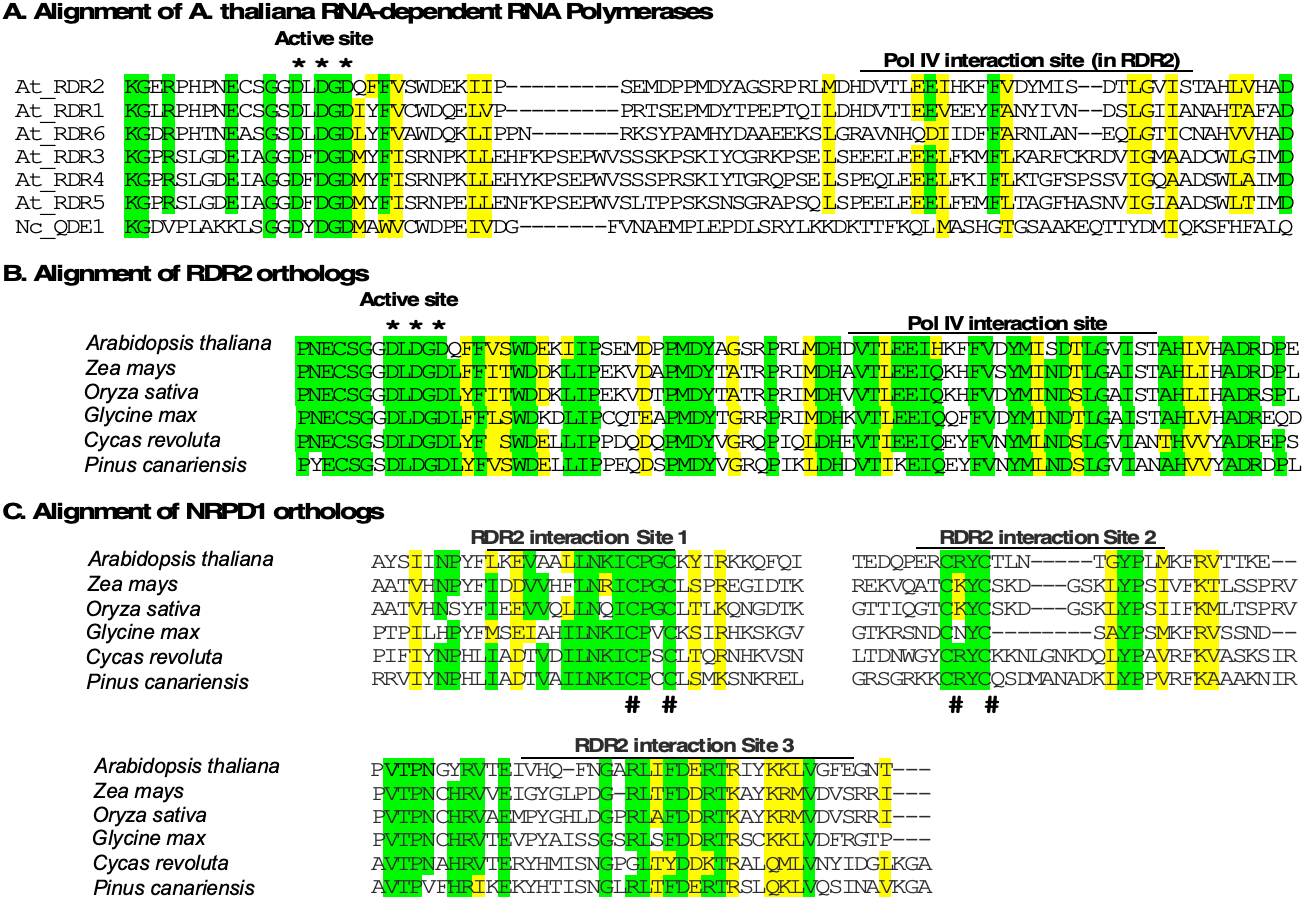
Sequence conservation at RDR2 and NRPD1 interaction sites. **A**. Alignment of *Arabidopsis thaliana* RNA-dependent RNA polymerases RDR1 through RDR6 and *Neurospora crassa* QDE-1, in the region that includes the conserved active site aspartates (*) and Pol IV (NRPD1) interaction site. Identical amino acids are highlighted in green, similar amino acids are highlighted in yellow. **B**. Sequence conservation at the Pol IV (NRPD1) interaction site among RDR2 orthologs of *A. thaliana*, maize (*Zea mays*), rice (*Oryza sativa*), soybean (*Glycine max*), cycad (*Cycas revoluta*) and pine (*Pinus canariensis*). **C**. Sequence conservation at the RDR2-interactions sites of NRPD1 orthologs of maize, rice, soybean, cycad, and pine.

RDR2 and NRPD1 co-purify in *A. thaliana*, a dicotyledenous (dicot) plant and maize, a monocotyledonous plant, whose last common ancestor existed approximately 150-200 million years ago. Figure 6B compares the NRPD1-interaction site of RDR2 and its orthologs in two dicots (Arabidopsis and Glycine), two monocots (Zea and Oryza) and two gymnosperms, Pinus and Cycas, who last shared a common ancestor with monocots and dicots ~400 million years ago. Substantial sequence conservation is apparent. Likewise, alignment of the three RDR2 interaction sites of Arabidiopsis NRPD1 with orthologs of the other species also reveals blocks of identical or similar amino acids (Figure 6C). Collectively, the high degree of sequence conservation at NRPD1 and RDR2 interactions sites suggest that Pol IV-RDR2 complexes may assemble in the same way throughout the plant kingdom.

## Discussion

Our results show that Pol IV and RDR2 form a stable complex via interactions between RDR2 and the largest of Pol IV’s 12 subunits. The three RDR2-interacting regions of NRPD1 are present within the first 300 amino acids of the protein. This region includes conserved domains A and B, which are two of eight domains (A-H) conserved among multisubunit RNA polymerase largest subunits, and ends just prior to the beginning of conserved domain C (see Figure 7 for positions of these domains). These first 300 amino acids include conserved and polymerase-specific elements of the structural module known as the clamp. Structural studies of bacterial, archaeal and eukaryotic RNA polymerases have shown that the clamp adopts different conformations, opening to allow DNA entry into the cleft where the catalytic site is located, and rotating inward to enclose the DNA-RNA hybrid that forms during transcription elongation (32–37). The clamp includes core and head subdomains, with clamp cores of eukaryotic Pols I, II, and III and archaeal RNA polymerases adopting nearly identical structures. By contrast, head subdomains are not conserved and form structures that facilitate interactions with polymerase-specific transcription factors or subunits. Smaller functional elements of the clamp include the zipper, lid and rudder loops that differ in sequence among different polymerases but are strategically located to help define the upstream end of the DNA-RNA hybrid and separate the RNA transcript from the DNA template at the base of the RNA exit channel (32–34). In Figure 7, the locations of these elements are shown based on their positions in Pol II (32, 33).

**Figure 7.**
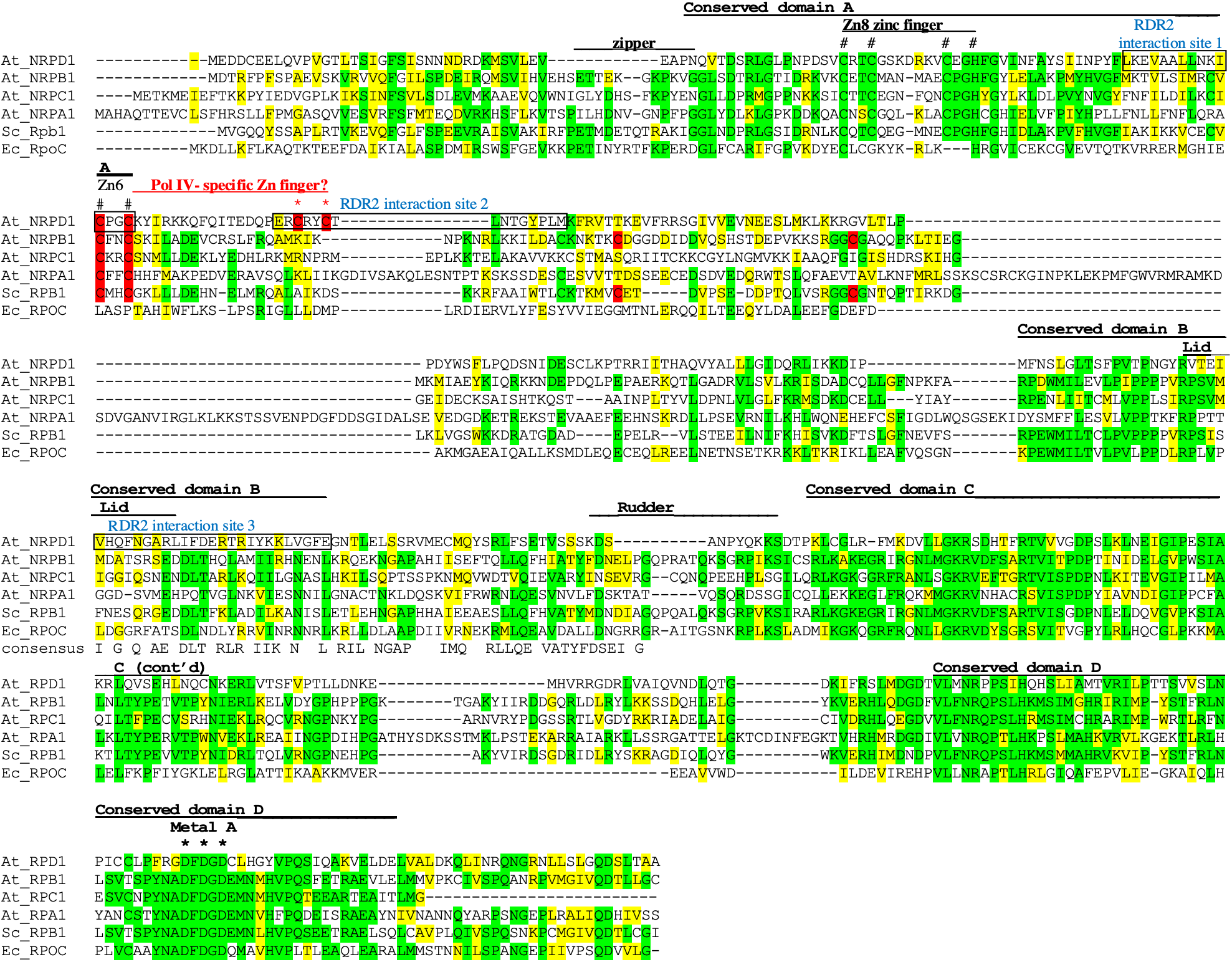
Comparison of amino terminal regions of RNA polymerase largest subunits. Multiple alignment of NRPD1’s amino terminus with corresponding sequences of the largest subunits of Arabidopsis (At) Pols I (NRPA1), II (NRPB1) and III (NRPC1), budding yeast Pol II (Sc_RPB1) and *E. coli* polymerase (Ec_RpoC). Identical amino acids in three or more sequences are shaded green; similar amino acids are shaded yellow. Positions of conserved domains A-D, zipper, lid and rudder elements are indicated based on their positions in yeast Pol II. The position of the Metal A site is also indicated, with black asterisks denoting the three magnesium-binding aspartates. Conserved Zinc coordinating amino acids of Zn8 and Zn6 are indicated with hash symbols (#). Cysteines of Zn6 are further highlighted in red, with cysteines of the Pol IV-specific CxxC, motif downstream of the conserved CxxC motif, indicated with red asterisks. RDR2 interacting regions 1, 2, and 3 of NRPD1, which overlap both Zn6 CxxC motifs and the lid element, respectively, are indicated in blue.

Within NRPD1, RDR2 interaction site 1 is located within conserved domain A, just downstream of the CxxC-CxxH zinc finger known as Zn8 (in Pol II) and including the first CxxC (highlighted in red and marked by hash symbols, #, in Figure 7) of the adjacent zinc coordinating element, Zn6. These zinc binding amino acids are invariant in the largest subunits of eukaryotic RNA polymerases I, II, III and IV (Figure 7). Genetic experiments have shown that substitutions at these invariant amino acids in Pol II confer temperature sensitive or lethal phenotypes (38). In Pols I, II and III, Zn8 and all preceeding sequences contribute to their structurally similar clamp cores whereas the conserved CxxC motif of Zn 6 contributes to their clamp heads. Given the conservation at these sites, it is likely that the same is true for Pol IV.

In yeast Pol II, Zn6 coordination involves the highly conserved CxxC motif and two cysteines (also highlighted in red in Figure 7) located 38 and 57 amino acids downstream, respectively. However, in NRPD1 the highly conserved Zn6 CxxC motif is followed, 17-20 amino acids downstream, by a second CxxC motif (highlighted in red and marked by asterisks in Figure 7) that is not found in the largest subunits of Pols I, II, III. This second, Pol IV-specific CxxC motif is part of RDR2-interaction site 2. Thus, RDR2 interaction sites 1 and 2 each include CxxC motifs, and both motifs are highly conserved (Figure 6C). These CxxC motifs in NRPD1 have the potential to coordinate a zinc atom, thereby generating a Pol IV-specific zinc finger within the RDR2 interacting region. However, our data suggest that formation of this putative zinc finger is not critical for RDR2 docking given that peptides that include the individual CxxC motifs (peptides 10 and 12) bind RDR2 in the peptide array experiment, as did flanking peptides, 9 and 13, which lacked the CxxC motifs. These results suggest that RDR2 can bind independently to either side of the putative zinc finger in a sequence-specific manner, consistent with the conservation of sequences adjacent to the CxxC motifs (see Figure 6C). Moreover, a tyrosine substitution at the first cysteine of the CxxC motif at the beginning of RDR2 interaction site 2 (C118Y, defining mutant allele *nrpd1-50*) did not disrupt RDR2 interaction, despite severly impairing Pol IV activity *in vivo* and *in vitro* (39). Thus the putative zinc finger may play a role in Pol IV transcription independent of RDR2 interaction.

NRPD1 has amino acid substitutions at the position of the lid (Figure 7) which in other polymerases is implicated in helping separate the RNA transcript from the DNA template at the upstream edge of the DNA-RNA hybrid. Interestingly, the amino acids at the expected position of the the lid overlap with RDR2 interaction site 3. Collectively the evidence that RDR2 binds structural elements of the clamp and lid suggests that RDR2 is located in proximity to the DNA/RNA hybrid and Pol IV’s RNA exit channel. This is consistent with RDR2’s use of Pol IV transcripts as templates and the coupling of Pol IV termination with RDR2 initiation (16).

At present, we lack high resolution structures for NRPD1 or RDR2. However, homology-based models in which NRPD1 sequences are fitted to the structure of yeast RPB1, and in which RDR2 sequences are fitted to the structure of Neurospora QDE1 (Figure 8), offer insights. The RDR2-interacting peptide regions of NRPD1 (highlighted in red) include helices of the clamp as well as the lid loop. In RDR2, the NRPD1 interaction site is a surface adjacent to the active site. The distance between the RDR2 interacting sites and the RNA strand of the DNA/RNA hybrid within NRPD1 and the distance between the NRPD1 interaction site and active site of RDR2 are similar (~22-25 Angstroms). Thus, if one imagines laying the RDR2 structure onto the NRPD1 structure, as if closing a book that has the images of the two proteins on facing pages, the RDR2 cleft has the potential to be located just above DNA-RNA hybrid and RNA exit channel of NRPD1. If so, this may account for RDR2’s ability to capture the released 3’ end of a terminated Pol IV transcript and explain the mechanistic coupling of Pol IV termination with RDR2 initiation. Structural studies of Pol IV, RDR2 and the complex formed by both proteins are now needed to test this hypothesis and further define the basis for Pol IV-RDR2 transcription coupling.

**Figure 8.**
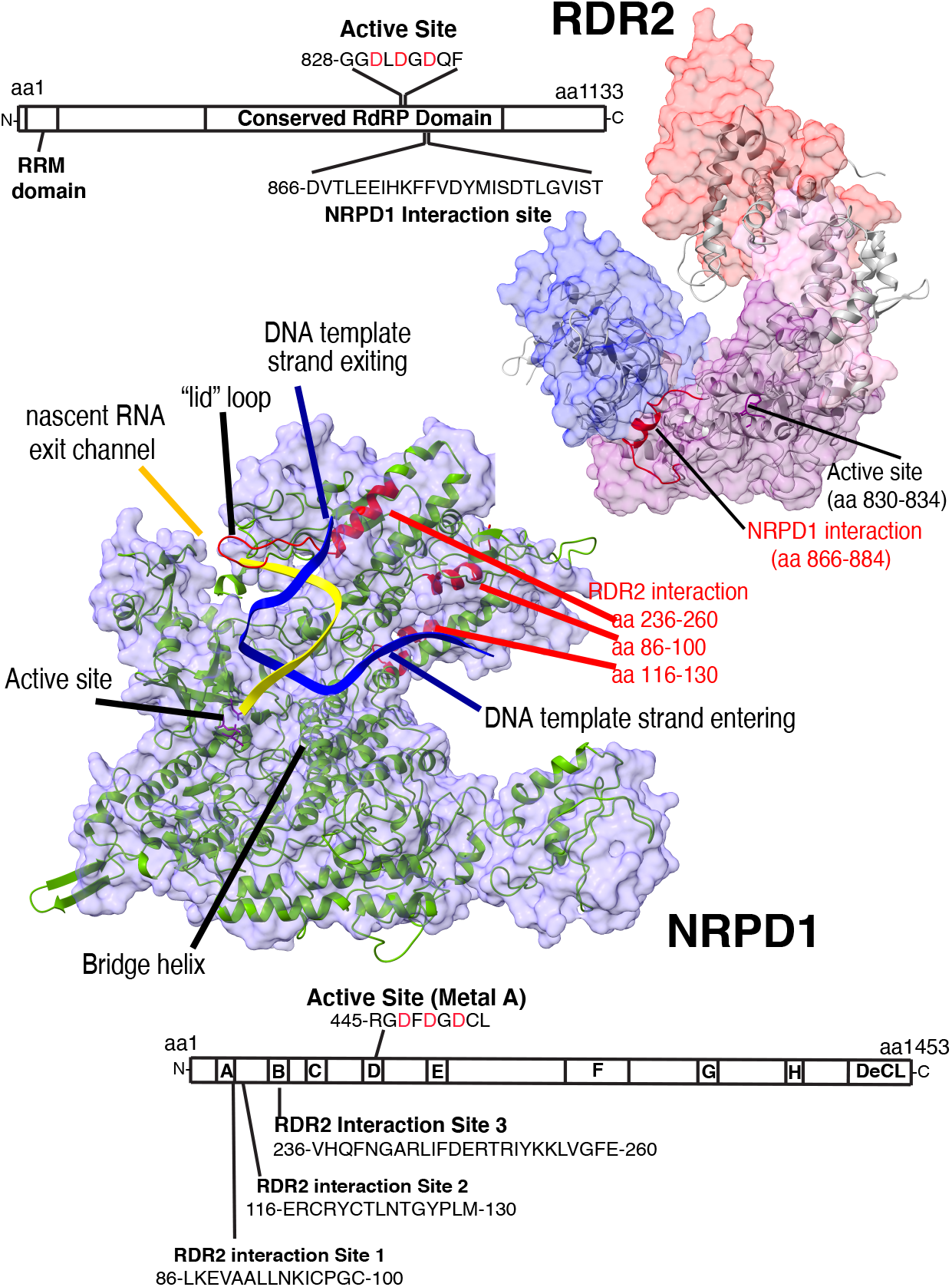
Structural homology models for RDR2 and NRPD1 showing predicted positions of their interaction sites. Models were generated using Phyre2 (Kelley et al., 2015) in intensive mode, integrating structural information from multiple homologs (see methods). At top right, the RDR2 structural model is shown as a gray ribbon superimposed onto a space-filling model for *Neurospora crassa* QDE-1 (31) whose slab, catalytic, neck and head domains are colored blue, purple, pink and red, respectively. Regions of RDR2 predicted to be disordered (amino acid interval 1-285 and 1122-1133) are excluded from the model. NRPD1-interacting sequences, aa 866-884, are highlighted in red. The diagram shows the relative locations of the active site and NRPD1-interaction site in a linear depiction of the RDR2 protein. At lower left, the NRPD1 structural model, shown as a green ribbon, was generated based on homologies to Pol II and Pol I and superimposed onto the space-filling model (purple) of the yeast Rpb1 subunit within a Pol II elongation complex (PDB:3HOV). The unique C-terminal region of NRPD1, aa 1276-1453, is excluded from the model. Template DNA and RNA are indicated as blue and yellow ribbons, respectively. Predicted positions of the three RDR2-interacting peptide motifs are highlighted. The diagram shows the relative locations of the active site and RDR2-interaction sites in a linear depiction of the NRPD1 protein. Positions of conserved domains A-H and of the DeCL domain, at the C terminus, are highlighted.

## Materials and Methods

### Cloning, expression, and purification of active or catalytically dead RDR2

An RDR2 cDNA expression construct encoding the wild-type protein fused to V5 epitope and 6xHis tags, was described previously (13). An active site mutant (RDR2-asm) was generated by site-directed mutagenesis using primers listed in Supplementary Table 1. The RDR2 and RDR2-asm genes were expressed in insect cells as BaculoDirect™ expression vectors (Thermo Fisher Scientific). Recombinant proteins were affinity purified using nickel-nitrilotriacetic acid (Ni-NTA) agarose (Qiagen) and elution with imidazole, followed by FPLC using Heparin-Sepharose and Superdex 200 columns. Purity was assessed by SDS-PAGE and staining with Coomassie Blue. See Supplemental information for additional details.

### Analytical size-exclusion chromatography

To estimate RDR2’s mass in solution, RDR2 and protein standards (GE Healthcare) were subjected to FPLC using a Superdex 200 10/300 GL column at 4°C using 50 mM HEPES-KOH pH7.5, 150 mM NaCl as the running buffer and a flow rate of 0.3 ml/min). Protein peaks were detected by UV absorbance at 280 nm, and SDS-PAGE with staining by Coomassie Blue.

### RDR2 activity assay

Conversion of single-stranded (ss) RNA into dsRNA was carried out in 50 μl reactions containing 200 ng RDR2, 25 nM template RNA (a 37 nt RNA 5’ end-labeled using T4 kinase and gamma-^32^P-ATP), 25 mM HEPES-KOH pH 7.5, 2 mM MgCl_2_, 0.1 mM EDTA, 0.1% Triton X100, 20 mM ammonium acetate, 3% PEG 8000, 0.1 mM each of ATP, GTP, CTP and UTP and 0.8 U/μL RiboLock RNase inhibitor (Thermo Fisher Scientific). Reactions were incubated at room temperature and stopped by adding EDTA to 10 mM. For RNase treatments, 1 unit of RNase ONE™ (Promega) or 0.01 unit of RNase V1 (Thermo Fischer Scientific) was added to reaction products and incubated for 15 minutes at 37 °C. Reactions were stopped by adding SDS to 0.1% (w/v). Reaction products were subjected to TRIzol extraction, precipitated with ethanol, resuspended in nuclease-free water, adjusted to 1X DNA loading dye (Thermo Fisher Scientific), and subjected to non-denaturing gel electrophoresis using a 15% polyacrylamide gel and 0.5X TBE (Tris-Borate-EDTA) running buffer. The gel was transferred to blotting paper, covered with plastic wrap and exposed to X-ray film at −80°C for 1 hour.

### Reconstitution of Pol IV-RDR2 complexes

Pol IV, Pol V, and Pol II were affinity-purified by virtue of their FLAG-tagged NRPD1, NRPE1 or NRPB2 subunits as described previously (29). To reconstitute Pol IV-RDR2 complexes, Pol IV expressed in an *rdr2* null mutant background was immobilized on anti-FLAG M2 resin then incubated with 1 μM recombinant RDR2 for 30 mins at 4 °C in 50 mM HEPES-KOH pH7.5, 150 mM NaCl. The resin was then washed three times with 100 μl of the same buffer to remove unbound proteins then subjected to SDS-PAGE and immunoblotting using anti-FLAG-HRP (Sigma Aldrich) or antibodies recognizing native NRPD1, NRPE1, or NRPB2. Anti-V5-HRP antibody (Invitrogen) was used to identify recombinant RDR2.

### Assay for dsRNA synthesis by Pol IV-RDR2 coupled reactions

Pol IV isolated in association with native RDR2, or Pol IV isolated from an *rdr2* null mutant (29) and reconstituted with recombinant RDR2, were tested for dsRNA synthetic ability as in Singh et al. (16). Reactions included 50 μl of anti-FLAG resin bearing immobilized Pol IV, with or without associated RDR2, in a total volume of 100 μL. Final concentrations of reaction components were 20 mM HEPES-KOH pH 7.6, 100 mM potassium acetate, 60 mM ammonium sulfate, 10 mM magnesium sulfate, 10% v/v glycerol, 10 μM zinc sulfate, 0.1mM PMSF, 1mM DTT, 0.8U/μL Ribolock™ (Thermo Fisher Scientific). To specifically label RNA strands synthesized by RDR2, reactions were performed using 250 nM T-less template DNA (lacking thymidines.), 1 mM each of GTP, CTP and UTP, 40 μM ATP and 10 μCi of [α^32^P]-ATP (3000 Ci/mmol; Perkin Elmer). Transcription reactions were incubated 1h at room temperature on a rotating mixer, stopped by addition of EDTA to 25 mM, and incubated at 75⁰ C for 10 min. Transcription reactions were then passed through PERFORMA spin columns (Edge Bio) and adjusted to 0.3 M sodium acetate (pH 5.2). 15 μg Glycoblue™ (Thermo Fisher Scientific) was added, and RNAs precipitated with 3 volumes of isopropanol at −20°C. Following centrifugation, pellets were washed with 70% ethanol and resuspended in 5 μl water. 5 μl of 2X RNA loading dye (New England Biolabs) was added, and the samples heated 5 min at 75⁰ C. RNAs were resolved on 15% polyacrylamide, 7M urea gels. Gels were transferred to filter paper, vacuum dried, and subjected to autoradiography or phosphorimaging.

### Cloning, expression, and purification of NRPD1

The NPRD1 open reading frame, fused to a C-terminal FLAG tag and codon optimized for insect cell expression, was synthesized by GenScript^®^and cloned into the baculovirus expression vector pKL-10xHis-MBP-SED-3C. The vector was first transfected into Sf9 insect cells to produce recombinant baculovirus particles, then High Five™ cells for NRPD1 over-expression. The NRPD1-Maltose Binding Protein (MBP) fusion protein was affinity purified using Amylose resin (New England Biolabs) and eluted with 20 mM maltose. The fusion protein was next subjected to anti-FLAG affinity purification, with in-column Prescission Protease digestion used to cleave the linker between MBP and NRPD1. Following extensive washing to remove free MBP, NRPD1 was eluted using an excess of FLAG peptide. See Supplemental information for a more detailed protocol.

### Cloning, over-expression and purification of NRPD1_1-300_ and RDR2_771-970_

NRPD1 amino acids 1-300 fused to His and FLAG tags, and RDR2 amino acids 771-971 fused to His and HA tags, were expressed in *E coli* using pET28 vectors. The proteins were then purified using nickel-NTA column chromatography and elution with imidazole. Purity was assessed by SDS-PAGE and immunoblotting using anti-FLAG-HRP or anti-HA-HRP antibodies. See Supplemental information for additional details.

### Co-immunoprecipitation experiments

Recombinant RDR2 and NRPD1 (0.25 μM each) were mixed and incubated for 30 minutes in 50 mM HEPES-KOH pH 7.5, 150 mM NaCl (at room temperature, in a volume of 100 μL. 50 μL aliquots were then mixed with 50 μL of anti-FLAG or anti-V5 antibody resin. After 30 minutes at room temp, the resins were washed twice with 200 μL 50 mM HEPES-KOH (pH 7.5), 150 mM NaCl. Proteins were eluted by addition of 6x SDS loading dye, and heating at 95°C for 2 minutes, then subjected to SDS-PAGE. Immunoblot analysis was used to detect the proteins using anti-FLAG-HRP to detect NRPD1 or anti-V5-HRP to detect RDR2 antibodies. The same methods were used to test for co-IP of full-length RDR2 with NRPD1_1-300_ or for co-IP of NRPD1_1-300_ with RDR2_771-970_.

### Yeast two-hybrid interaction assay

*Saccharomyces cerevisiae* strain MaV203 was transformed according to the Matchmaker protocol (Thermo Fisher) with plasmids expressing NRPD1 or RDR2 fused to the GAL4 activation domain or GAL4 DNA-binding domain. Transformants were selected on SC-Leu-Trp plates for 3 days at 30 °C. To test for interactions, each strain was replica plated onto SC,-Trp,-Leu and SC,-Trp,-Leu,-His, +3AT agar. Growth after 3 days at 30 °C was then assessed.

### Expression of NRPD1 and RDR2 polypeptides

20-33 kD subregions of NRPD1 and RDR2 polypeptides were expressed *in vitro* using a PURExpress^®^ *In Vitro* Protein Synthesis Kit (New England Biolabs). Briefly, the desired intervals of the synthetic genes were amplified using the primers in Supplementary Table 1 and then used as templates for *in vitro* transcription-translation reactions.

### Peptide array protein interaction assays

Custom peptide libraries corresponding to NRPD1 amino acids 1-300 or RDR2 amino acids 771-971 were obtained from Genscript and dot-blotted (10 ng) onto nitrocellulose. Filters were submerged in 5% (w/v) nonfat dry milk in 1X TBST (Tris buffered saline, 0.1 % Tween 20), 30 min at room temperature to block free protein binding sites. The blocking solution was then discarded. NRPD1 peptide arrays were then probed with ~200 ng of RDR2_771-971_-HA in 8 ml of 50 mM HEPES-KOH pH 7.5, 100 mM NaCl at room temperature for 1 hour. The RDR2 peptide array was probed with ~200 ng of NRPD1_1-300_-FLAG. Filters were then washed twice with 8 ml of 1X TBST, then incubated with anti-FLAG-HRP or anti-HA-HRP in 8 ml 1X TBST at room temperature for 1 hour. The filters were washed twice with 8 ml 1X TBST, then developed using a Pierce™ Enhanced Chemiluminescence Western Blotting Kit (Thermo Fisher Scientific).

### Crosslinking-Mass Spectroscopy

Purified RDR2_771-970_ and NRPD1_1-300_ proteins were pre-incubated then crosslinked using 0.1 mM Bissulfosuccinimidyl suberate (BS3). Crosslinked protein complexes were reduced using 10 mM TCEP (Tris(2-carboxyethyl)phosphine), alkylated with 20 mM iodoacetamide, and digested with 12.5 ng/μL each of Trypsin and Chymotrypsin for 16 hr. Resulting peptides were resolved by HPLC using a C18 column using an acetonitril gradient and subjected to electrospray ionization and anlyzed using an Orbitrap Fusion Lumos mass spectrometer. Data are available at the ProteomeXchange Consortium via the PRIDE partner repository (40) with the dataset identifier PXD020170. See Supplemental information for a detailed protocol.

### Molecular modeling of NRPD1 and RDR2

A structural model for NRPD1 was generated using Phyre2 protein structure prediction software (41), in intensive mode, based on homologies to six Pol II or Pol I largest subunits. The RDR2 model is based on two fungal QDE-1 RNA-dependent RNA polymerases. Resulting models were then superimposed onto the structures of budding yeast Rpb1 (the Pol II largest subunit; PDB:3HOV) (42) or *Neurospora crassa* QDE-1 (PDB:2J7N) (31) crystal structures using the MatchMaker tool of Chimera (43). Superimposed models were then imported to ChimeraX (44) and solvent-excluded surfaces of RPB1 and QDE-1 were visualized and colored. See Supplemental information for additional details.

### Sequence alignments

Amino acid sequence alignments were performed using CLUSTAL W. Conserved sequences were highlighted using BOXSHADE V3.31.

## Acknowledgments

VM dedicates this work in loving memory of his father, Dr. Umesh Chandra Mishra. We thank the Drosophila Genome Resource Center at Indiana University Bloomington for access to their insect cell culture facilities. We thank Tsuyoshi Imasaki for his efforts in engineering the baculovirus vector used for NRPD1 overexpression.

## Author Contributions

JS performed the Pol IV-RDR2 coupled transcription assay and generated the NRPD1 recombinant baculovirus, using a vector engineered by YT. AF purified NRPD1. FW cloned NRPD1 and conducted the yeast assays. YZ and JCT performed mass spectroscopy and aidentified cross-linked peptides. VM performed all other experiments. CSP and VM wrote the manuscript.

