## Supplemental Information for "Assembly of a double-stranded RNA synthesizing complex: RNA-DEPENDENT RNA POLYMERASE 2 docks with NUCLEAR RNA POLYMERASE IV at the clamp domain"

### Detailed methods

#### Cloning, expression, and purification of active or catalytically dead RDR2

A recombinant RDR2 gene encoding the wild-type open reading frame fused to a V5 epitope tag and 6xHis tag, was described previously (1). An active site mutant (RDR2-asm), bearing alanine substitutions at three amino acids of the magnesium ion binding site, was generated from the wild-type RDR2 gene using a Q5® Site-Directed Mutagenesis Kit (New England Biolabs Inc.) and primers listed in Supplementary Table 1.

Recombinant RDR2 and RDR2-asm genes were first cloned into pENTR plasmids then linearized and recombined into the BaculoDirect™ expression vector (Thermo Fisher Scientific) using LR clonase reactions. Resulting constructs were transfected into Sf9 insect cells and passaged to increase viral titer according to the manufacturer's instructions (Thermo Fisher Scientific). High Five™ *Trichoplusia ni* cells (Thermo Fisher Scientific) were then infected for protein overexpression experiments. Cells were lysed in 50 mM HEPES-KOH (pH 7.5), 400 mM KCl, 1 mM PMSF, 5 mM benzamidine-HCl, and 10% glycerol. Lysates were then subjected to centrifugation at 39,000 x g for 30 min at 4°C. The supernatant was then loaded onto a nickel-nitrilotriacetic acid (Ni-NTA) agarose (Qiagen) column pre-equilibrated in 50 mM HEPES-KOH pH 7.5 and 400 mM KCl. The column was subsequently washed in three steps with buffer containing 50 mM HEPES-KOH pH 7.5, 300 mM NaCl and 5, 10 or 15 mM imidazole, respectively. Proteins still bound to the column were then eluted using buffer containing 400 mM imidazole. Eluted proteins were dialyzed into 50 mM HEPES-KOH pH 7.5, 50 mM NaCl and applied to a Heparin-Sepharose column equilibrated in the same buffer. After washing using the dialysis/column equilibration buffer, bound RDR2 was eluted with 50 mM HEPES-KOH pH 7.5,

300 mM NaCl. The eluted fraction was concentrated to a final volume of 1 ml using a centrifugal spin filter (Millipore) with a 30 KDa cutoff size. The concentrated sample was then loaded onto a Superdex 200 10/300 GL column connected to an AKTA FPLC instrument (GE Healthcare) and subjected to gel filtration chromatography using 50 mM HEPES-KOH pH7.5, 150 mM NaCl as the running buffer. Peak fractions were pooled, dialyzed into 50 mM HEPES-KOH pH 7.5, 150 mM NaCl, and 50% glycerol and stored at -20 °C. To assess purity, samples of peak fractions were subjected to SDS-PAGE on a 4-16 % gradient gel and proteins were visualized by staining with Coomassie Blue.

**Cloning, expression, and purification of NRPD1**

A synthetic NPRD1 open reading frame, in-frame with a C-terminal FLAG tag and codon optimized for insect cell expression, was synthesized by GenScript® and sub-cloned into the *Bam*HI and *Hind*III sites of the baculovirus expression vector pKL-10xHis-MBP-SED-3C, yielding the expression construct, pKL-10xHis-MBP-SED-3C-NPRD1-FLAG (pYT1107). Detailed description of the pKL-10xHis-MBP-SED-3C vector, and its utility for expression of large proteins, will be described elsewhere (Imasaki and Takagi, manuscript in preparation), but briefly, the vector was engineered by inserting a DNA cassette encoding a 10x His tag and maltose-binding protein (MBP) followed by a synthetic protein, termed SED, and a human rhinovirus 3C protease (HRV 3C Protease) site (LEVLFQGP) immediately 5' of the *Bam*HI site of the pKL baculovirus expression vector (2) using the SLIC method (3). The DNA cassette was synthesized by GenScript (Piscataway, NJ), with protein-coding regions codon-optimized for insect cell expression.

The pKL-10xHis-MBP-SED-3C-NPRD1-FLAG expression vector was transformed into DH10Bac competent cells (Thermo Fisher Scientific) and plated on LB agar containing X-gal and IPTG, allowing recombinant NRPD1 transformants to be selected by blue-white screening. DNA was isolated from the positive clones using PureLink viral DNA purification kit (Thermo Fisher Scientific) and transfected into Sf9 insect cells to produce recombinant baculovirus particles. High Five™ cells were then infected, at a multiplicity of infection of 1, for NRPD1 over-expression using 500 ml cell suspension cultures. Cell pellets were frozen, then resuspended in 125 ml lysis buffer containing 50 mM HEPES pH7.9, 400 mM KCl, 10% glycerol, 5 mM DTT, 1 mM PMSF, 1x Protease Inhibitor Cocktail (Sigma, P9599) and incubated on ice for 30 min, with occasional mixing. The lysate was subjected to centrifugation at 153,720 x g for 45 min at 4˚C, and the supernatant passed through a 0.22 $\mu$m syringe filter. Amylose resin (New England Biolabs) equilibrated in lysis buffer was added as a slurry, equivalent to a packed bed volume of 1 ml, to the clarified lysate and incubated, with rotation, at 4°C for 2 hours. After low-speed centrifugation to pellet the resin, the supernatant was removed and discarded. The amylose resin, with associated proteins, was then transferred to an open column and washed, by gravity flow, with 15 column volumes of lysis buffer, followed by 3 column volumes of low-salt buffer [50 mM HEPES pH7.9, 100 mM potassium acetate, 10% glycerol, 5 mM DTT, 0.2 mM PMSF]. Proteins were eluted from the resin using 4 column volumes of elution buffer [50 mM HEPES pH7.9, 300 mM potassium acetate, 20 mM maltose, 10% glycerol, 5 mM DTT, 0.2 mM PMSF]. 200 $\mu$l of M2 anti-FLAG affinity gel (A2220, Sigma) was the added to the eluted fraction and incubated, with rotation at 4 °C, for 3 hours. The anti-FLAG resin, and associated NRPD1, was collected by centrifugation, and the supernatant removed. The resin was washed three times, 15 min each, at 4°C in 2 ml high salt buffer [50 mM Tris-HCl pH7.9, 400 mM NaCl,

1 mM DTT, 10% glycerol, 0.2 mM PMSF] using a rotating mixer. The anti-FLAG resin was then equilibrated in protease reaction buffer [50 mM Tris-HCl pH7.9, 150 mM NaCl, 1 mM DTT, 10% glycerol, 0.2 mM PMSF] such that the resin accounted for 50% of the total volume. 1/40 volume of Prescission Protease (Human Rhinovirus 3C protease; GE Healthcare) was then added to cleave away N-terminal region that includes the Maltose Binding Protein from the recombinant NRPD1 protein. After overnight digestion at 4 °C, the resin was pelleted by centrifugation at 300 x g, 2 min and the supernatant removed. The anti-FLAG resin with bound NRPD1 was transferred to an open gravity-flow column and washed three times with 10 column volumes of ATP wash buffer [50 mM HEPES pH7.9, 400 mM NaCl, 5 mM ATP-magnesium salt, 1 mM DTT, 10% glycerol, 0.2 mM PMSF]. NRPD1 was then eluted from the resin using FLAG elution buffer [50 mM Tris-HCl pH7.9, 150 mM NaCl, 1 mM DTT, 10% glycerol, 0.2 mM PMSF, 0.5 mg/ml 3xFLAG peptide (APExBIO)]. An Amicon Ultracel-30K Ultra Centrifugal Filter (Merck Millipore) was then used to concentrate the purified recombinant NRPD1.

**Cloning, over-expression and purification of $NRPD1_{1\text{-}300}$ and $RDR2_{771\text{-}970}$**

NRPD1 amino acids 1-300 fused to His and FLAG tags, and RDR2 amino acids 771-971 fused to His and HA tags, were cloned into pET28a and expressed in *E coli* BL-21 cells upon induction using 1 mM of isopropyl-1-thio-β-D-galactopyranoside (IPTG). An 800 ml suspension culture, grown at 16 °C, was used for protein overexpression. Cells were harvested by centrifugation at 3800 x g for 10 mins, 4˚C and the pellets were resuspended in 50 ml 50 mM HEPES-KOH pH 7.5, 300 mM NaCl, 10% glycerol and 1 mM PMSF. Resuspended cells were lysed by sonication, using a probe sonicator, and the lysates were clarified by centrifugation at 22,000 x g for 30 mins at 4 °C. To affinity purify the proteins based on their His tags, the supernatants were applied to nickel-NTA resin columns pre-equilibrated with 50 mM HEPES-KOH pH 7.5 and 300 mM NaCl. The columns were washed successively with lysis buffer containing 20, and then 40 mM imidazole, and remaining proteins were then eluted with lysis buffer containing 400 mM imidazole. The purity of the eluted recombinant proteins was assessed by SDS-PAGE and Coomassie Blue staining as well as immunoblot analysis using anti-FLAG-HRP antibody (Sigma-Aldrich) to detect $NRPD1_{1\text{-}300}$ or anti-HA-HRP antibody (Sigma-Aldrich) to detect $RDR2_{771\text{-}970}$.

**Crosslinking-Mass Spectroscopy**

Purified $RDR2_{771\text{-}970}$ and $NRPD1_{1\text{-}300}$ proteins were pre-incubated then mixed with 0.1 mM Bissulfosuccinimidyl suberate (BS3) crosslinker (Thermo Fisher Scientific) in 50 mM HEPES-KOH pH7.5, 150 mM NaCl buffer, in a total volume of 100 ml, and incubated at room temperature for 30 min. The reaction was quenched by adding 5 µl of 1 M ammonium bicarbonate for 5 min. Following SDS-PAGE, gel regions corresponding to the masses of crosslinked protein complexes were diced with a razor blade and reduced for 45 min at 56 °C in 100 ml of 10 mM TCEP (Tris(2-carboxyethyl)phosphine) followed by alkylation with 20 mM iodoacetamide for 1 h in the dark at 21 °C. Proteins were then digested with 12.5 ng/µL each of Trypsin and Chymotrypsin for 16 hr. Resulting peptides were separated using an Easy-nLC 100 HPLC system with an Acclaim PepMap™ RSLC C18 analytical column (75 µm × 150 mm, 2 µm, 100 Å) using an acetonitrile-based gradient (Solvent A: 0% acetonitrile, 0.1% formic acid; Solvent B: 80% acetonitrile, 0.1% formic acid) at a flow rate of 300 nl/min. A 30 min gradient was as follows: 0-0.5 min, 2-8% B; 0.5-24 min, 8-40% B; 24-26 min, 40-100% B; 26-30 min,

100% B, followed by re-equilibration to 2% B. The electrospray ionization was carried out with a nano-ESI source at a 260 °C capillary temperature and 1.9 kV spray voltage. The Orbitrap Fusion Lumos mass spectrometer (Thermo Scientific, Bremen, Germany) was operated in data-dependent acquisition mode with mass range 400 to 2000 m/z. Precursor ions with charge state from 3 to 7 were selected for tandem mass (MS/MS) analysis with 3 sec cycle time using HCD at 30% collision energy. Intensity threshold was set at 2e5. The dynamic exclusion was set with a repeat count of 1 and exclusion duration of 30 s. The resulting data was searched with Protein Prospector (http://prospector.ucsf.edu/prospector/mshome.htm) against RDR2 (aa 771-970) and NRPD1 (aa 1-300). Carbamidomethylation of cysteine residues was set as a fixed modification. Protein N-terminal acetylation, oxidation of methionine, protein N-terminal methionine loss, pyroglutamine formation were set as variable modifications. Crosslinking search type was set as Disuccinimidyl suberate (DSS), a water-insoluble analog of BS3 which has the same mass addition. A total of three variable modifications were allowed. Trypsin and Chymotrypsin digestion specificity with two missed cleavage was allowed. The mass tolerance for precursor and fragment ions was set to 5 and 10 ppm, respectively.

Crosslinked peptides were manually validated and categorized into three groups. Each peptide in a crosslink received an individual score, and the high confidence group having at least three unique y or b-type fragment ions derived from the lower scoring peptide. The low confidence group included crosslinked peptides whose lower-scoring peptide had had only one or two unique fragment ions. The mass-alone group meant the lowest scoring crosslinked peptide had fewer than five amino acid residues, such that there is a much reduced chance of observing three unique fragment ions.

The mass spectrometry proteomics data have been deposited to the ProteomeXchange Consortium via the PRIDE partner repository (4) with the dataset identifier PXD020170.

**Molecular modeling of NRPD1 and RDR2**

NRPD1 and RDR2 structural models were generated using Phyre2 protein structure prediction software (5) used in intensive mode to integrate comparisons to multiple homologs with known structures. The NRPD1 (NP_001185298.1) model was generated based on homologies to six homologs: *Schizosacharomyces pombe* Rpb1 (the yeast Pol II largest subunit; PDB:3H0G, chain A) (6); *Sus scrofa* Rpb1 (the largest subunit of mammalian Pol II within a paused Pol II-DSIF-NELF transcription complex (PDB:6GML, chain A) (7); *Bos taurus* Rpb1 (the largest subunit of mammalian Pol II in a transcription elongation complex; PDB:5FLM, chain A) (8); *Saccharomyces cerevisiae* Rpb1 (the largest subunit of yeast Pol II in a transcription elongation complex; PDB:1TWF, chain A) (9); *Saccharomyces cerevisiae* Rpa1 (the yeast Pol I largest subunit; PDB:4C3I, chain A) (10); *Saccharomyces cerevisiae* Rpa1 (the largest subunit of yeast Pol I, in an elongation complex; PDB:5M3F, chain A) (11). For RDR2 (NP_192851.1, 1133aa), two fungal RNA-dependent RNA polymerases served as models for RDR2 amino acids 284-1122: *Neurospora crassa* QDE-1 (PDB :2J7N, chain A) (12) and *Thermothielavioides terrestris* QDE-1 (PDB : 5FSW, chain C) (13). The predicted models were then superimposed onto Saccharomyces cerevisiae Rpb1 (the Pol II largest subunit; PDB:3HOV, chain A) (14) or *Neurospora crassa* QDE-1 (PDB:2J7N, chain B) (12) crystal structures using the MatchMaker tool of Chimera (15). Superimposed models were then imported to ChimeraX (16) and solvent-excluded surfaces of RPB1 and QDE-1 were visualized and colored.

**Table S1. Primer sequences used for RDR2 and NRPD1 subcloning**

| | |
|---|---|
| RDR2 F* | CACCATGGTGTCAGAGACGACGACGAAC |
| RDR2 R** | TCAAATGGATACAAGTCCACTTGTTTT |
| RDR2 (DGD-AAA) F | AGCACAGTTTTTTGTTAGCTGGGATG |
| RDR2 (DGD-AAA) R | GCTGCGAGATCGCCACCAGAACA |
| NRPD1 F | GGATCCATGGAAGACGACTGCGAGGAA |
| NRPD1 R | AAGCTTTTAGCAGAGCTTAGTATCGAA |
| NRPD1 (1-300) F | GCGAATTAATACGACTCACTATAGGGCTTAAGTATAAGGAGGAAAAAATATGGAAGACGACTGCGAGGAACTGCAA |
| NRPD1 (1-300) R | AAACCCCTCCGTTTAGAGAGGGGTTATGCTAGTTACTTGTCGTCATCGTCTTTGTAGTCGGTATCGCTCTTCTTTTGGTACGG |
| NRPD1 (255-555) F | GCGAATTAATACGACTCACTATAGGGCTTAAGTATAAGGAGGAAAAAAT ATGAAGCTCGTGGGTTTCGAGGGCAACACA |
| NRPD1 (255-555) R | AAACCCCTCCGTTTAGAGAGGGGTTATGCTAGTTACTTGTCGTCATCGTCTTTGTAGTCGAAACCGGGCGGGAACAGCATACC |
| NRPD1 (482-781) F | GCGAATTAATACGACTCACTATAGGGCTTAAGTATAAGGAGGAAAAAAT ATGAACGGTAGAAACCTGCTCTCCTTGGGC |
| NRPD1 (482-781) R | AAACCCCTCCGTTTAGAGAGGGGTTATGCTAGTTACTTGTCGTCATCGTCTTTGTAGTCGCCCTTAGCTCCTCTCAGTGGGGA |
| NRPD1 (709-1009) F | GCGAATTAATACGACTCACTATAGGGCTTAAGTATAAGGAGGAAAAAAT ATGAGGGATGTCCAGGCTTTGGCCTACAGA |
| NRPD1 (709-1009) R | AAACCCCTCCGTTTAGAGAGGGGTTATGCTAGTTACTTGTCGTCATCGTCTTTGTAGTCGTATTGTTCGTTGAGCGATGAAAC |
| NRPD1 (935-1235) F | GCGAATTAATACGACTCACTATAGGGCTTAAGTATAAGGAGGAAAAAAT ATGAAGAAGAAGCACGGATTCGAGTACGGT |
| NRPD1 (935-1235) R | AAACCCCTCCGTTTAGAGAGGGGTTATGCTAGTTACTTGTCGTCATCGTCTTTGTAGTCCTCCTTGGCAGCCTTCAGGAAACA |
| NRPD1 (1163-1453) F | GCGAATTAATACGACTCACTATAGGGCTTAAGTATAAGGAGGAAAAAAT ATGATCTTCGTGGCTAACCTCGAGTCGGCC |
| NRPD1 (1163-1453) R | AAACCCCTCCGTTTAGAGAGGGGTTATGCTAGTTACTTGTCGTCATCGTCTTTGTAGTCTGGGTTTTCTGAGAAGCCACCGCT |
| RDR2 (1-200) F | GCGAATTAATACGACTCACTATAGGGCTTAAGTATAAGGAGGAAAAAATATGGTGTCAGAGACGACGACGAACCGA |
| RDR2 (1-200) R | AAACCCCTCCGTTTAGAGAGGGGTTATGCTAGTTAAGCGTAATCTGGAACATCGTATGGGTAGGCTATGTGAACTGTTACTCTTTT |
| RDR2 (155-354) F | GCGAATTAATACGACTCACTATAGGGCTTAAGTATAAGGAGGAAAAAATATGAAGATTGAAGTTAGGTTTGAGGATATT |
| RDR2 (155-354) R | AAACCCCTCCGTTTAGAGAGGGGTTATGCTAGTTAAGCGTAATCTGGAACATCGTATGGGTACTGAGTCTTGACAAAGAAGACAGG |
| RDR2 (309-508) F | GCGAATTAATACGACTCACTATAGGGCTTAAGTATAAGGAGGAAAAAATATGAGTCTTTTTGCTGCGTCAGATATGGAA |
| RDR2 (309-508) R | AAACCCCTCCGTTTAGAGAGGGGTTATGCTAGTTAAGCGTAATCTGGAACATCGTATGGGTAGATTTTACGGAAACAACCCATCCA |
| RDR2 (463-662) F | GCGAATTAATACGACTCACTATAGGGCTTAAGTATAAGGAGGAAAAAATATGACCGTTGGGCCTAAGAGGTTTGAGTTC |
| RDR2 (463-662) R | AAACCCCTCCGTTTAGAGAGGGGTTATGCTAGTTAAGCGTAATCTGGAACATCGTATGGGTAATTCCCAAGCATAGATAAATGCAC |
| RDR2 (617-816) F | GCGAATTAATACGACTCACTATAGGGCTTAAGTATAAGGAGGAAAAAATATGCTGAACGTTACCAGGTGGACAGAG |
| RDR2 (617-816) R | AAACCCCTCCGTTTAGAGAGGGGTTATGCTAGTTAAGCGTAATCTGGAACATCGTATGGGTACTGAGGAAAGATGATGCAGTCGAG |
| RDR2 (771-970) F | GCGAATTAATACGACTCACTATAGGGCTTAAGTATAAGGAGGAAAAAATATGGAAACATCTGTGGTTATTGGGAAAGTG |
| RDR2 (771-970) R | AAACCCCTCCGTTTAGAGAGGGGTTATGCTAGTTAAGCGTAATCTGGAACATCGTATGGGTACTGTGCTAGAGAGCTCTTCACAGC |
| RDR2 (925-1133) F | GCGAATTAATACGACTCACTATAGGGCTTAAGTATAAGGAGGAAAAAATATGGGAGCTCCAGCTGAGATGCCTTATGCC |
| RDR2 (925-1133) R | AAACCCCTCCGTTTAGAGAGGGGTTATGCTAGTTAAGCGTAATCTGGAACATCGTATGGGTAAATGGATACAAGTCCACTTGTTTT |
| NRPD1 (1-300)-pET28a F | ATGGATCCATGGAAGACGACTGCGAGGAACTGCAA |
| NRPD1 (1-300)-pET28a R | CCAAGCTTTTACTTGTCGTCATCGTCTTTGTAGTCGGTATCGCTCTTCTTTTGGTA |
| RDR2 (771-970)-pET28a F | ATGGATCCATGGAAACATCTGTGGTTATTGGGAAA |
| RDR2 (771-970)-pET28a R | CCAAGCTTTTAAGCGTAATCTGGAACATCGTATGGGTACTGTGCTAGAGAGCTCTTCACAGC |

*F = forward primer    **R = Reverse primer

### Table S2. Oligonucleotides used for transcription assays

| | |
|---|---|
| RDR2 Transcription template | rUrArCrArArGrCrGrArArUrGrArGrUrCrArUrUrCrArUr CrCrUrArArGrUrCrCrArArCrArUrA |
| T-Less Template DNA strand | CAAAAACGAGACAGACAACGAAAGCAGACAGAGAACG CCAGGACCGACACG |
| Non-Template DNA strand | GCTGCTTTCGTTGTCTGTCTCGTTTTTG |
| RNA Primer (hybridizes to T-Less template) | rUrGrCrArUrArArArGrUrCrCrUrGrGrC |

**A. Expression of six overlapping subregions of NRPD1**

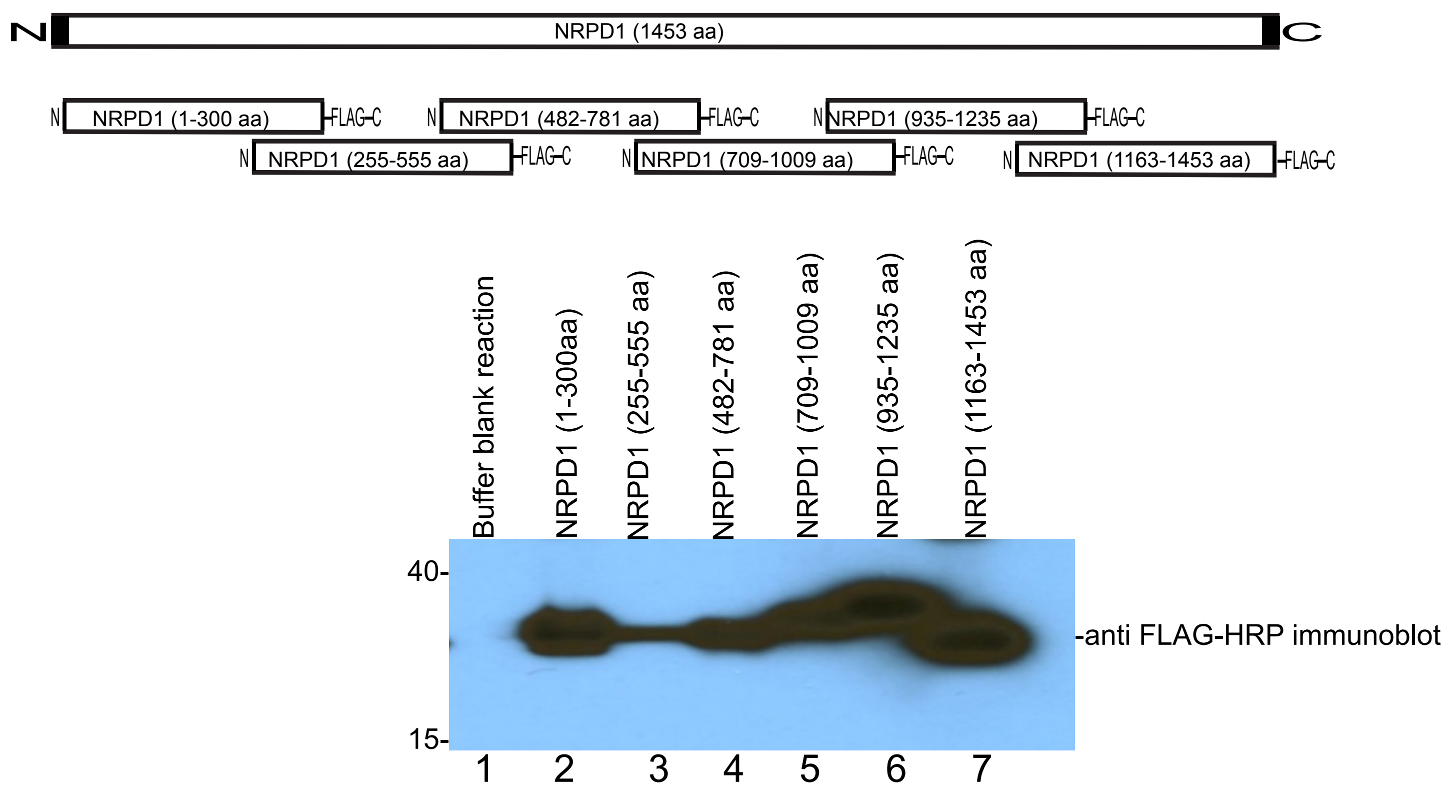


**B. Over-expression and purification of NRPD1$_{1-300}$**

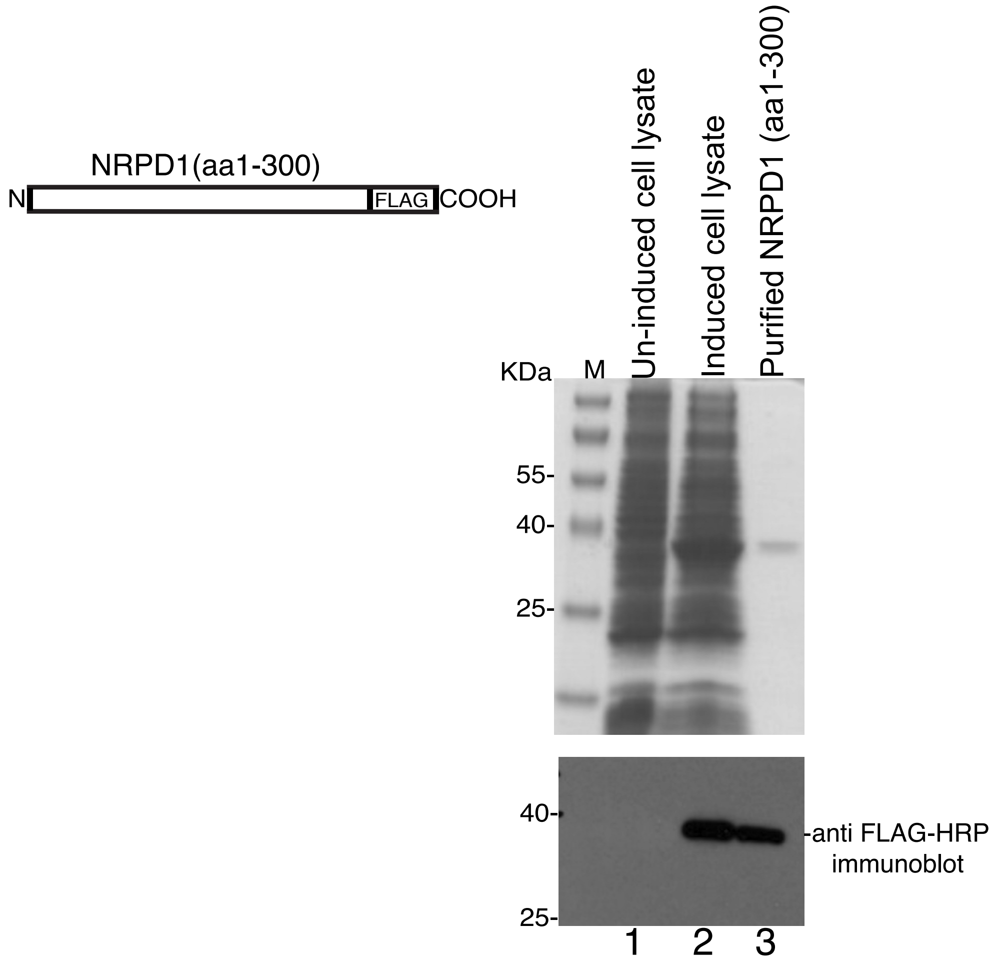


**Figure S1. Subdividing NRPD1 into subregions to test for RDR2 interaction.**
A. The diagram at top depicts six recombinant expression constructs, each encoding ~33 kDa polypeptides that collectively represent the 1453 amino acids of full-length NRPD1. The corresponding amino acid intervals of NRPD1 are indicated in parentheses. Each polypeptide was designed to have a FLAG epitope tag at the C-terminus. The polypeptides were expressed in vitro using a cell free transcription-translation system then subjected to SDS-PAGE and immunoblot detection using anti-FLAG antibodies conjugated to horseradish peroxidase (HRP).
B. The open reading frame for the amino-terminal segment of NRPD1, amino acids 1-300 was cloned into an E. coli expression plasmid and overexpressed in bacterial cells. The stained gel shows size markers (M), uninduced cells, cells treated with IPTG to induce expression of the poly-peptide, and purified NRPD1$_{1-300}$. The image at the bottom is an immunoblot of the same fractions probed with anti-FLAG antibody conjugated to horseradish peroxidase (HRP).

### A. Expression of seven overlapping subregions of RDR2

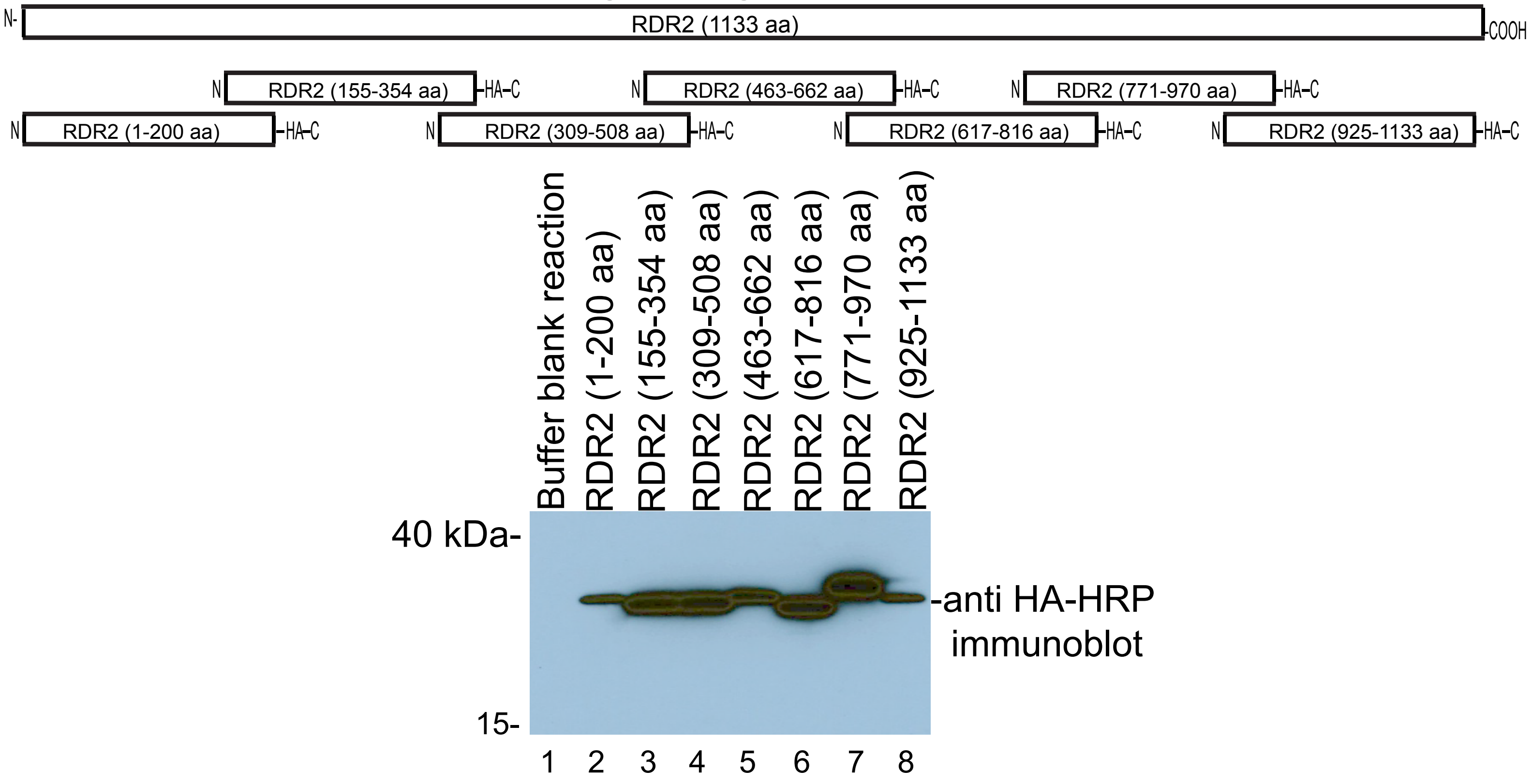


### B. Over-expression and purification of $RDR2_{771-970}$

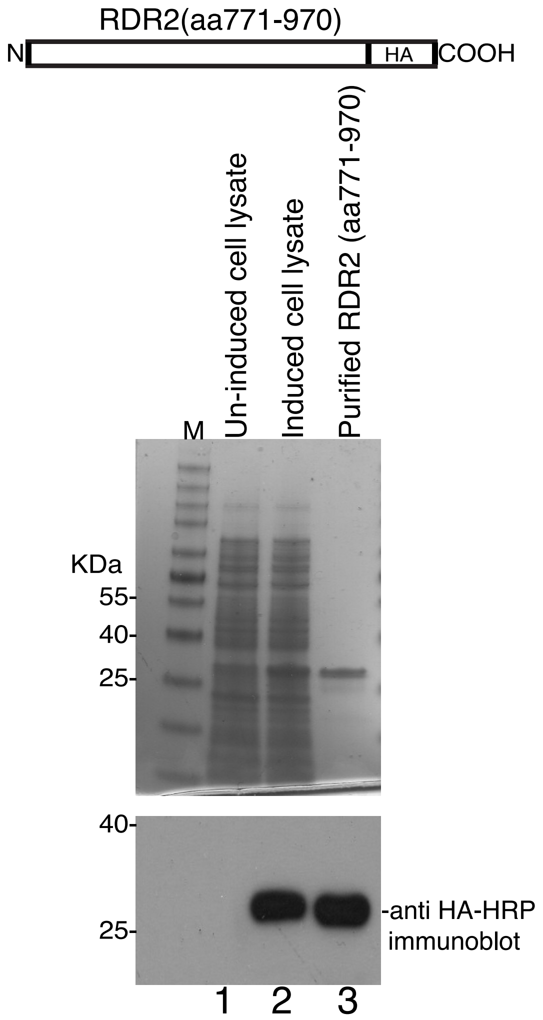


**Figure S2. Subdividing RDR2 into subregions to test for NRPD1 interaction.**
A. The diagram at top depicts seven recombinant expression constructs, each encoding ~25 kDa polypeptides that collectively represent the 1133 amino acids of full-length RDR2. The corresponding amino acid intervals of RDR2 are indicated in parentheses. Each polypeptide was designed to have a HA epitope tag at the C-terminus. The polypeptides were expressed in vitro using a cell free transcription-translation system then subjected to SDS-PAGE and immunoblot detection using anti-HA antibodies conjugated to horseradish peroxidase (HRP).
B. The open reading frame for the amino-terminal segment of NRPD1, amino acids 771-970 was cloned into an E. coli expression plasmid and overexpressed in bacterial cells. The stained gel shows size markers (M), uninduced cells, cells treated with IPTG to induce expression of the polypeptide, and purified $RDR2_{771-970}$. The image at the bottom is an immunoblot of the same fractions probed with anti-HA antibody conjugated to horseradish peroxidase (HRP).
